# A Dual-Locus-Targeting Strategy to Enhance CRISPR/Cas9-mediated CFTR Replacement via Helper-Dependent Adenoviral vector in porcine genome

**DOI:** 10.64898/2026.06.10.731381

**Authors:** Ziyan Rachel Chen, Zhichang Peter Zhou, Rongqi Cathleen Duan, Amy Wong, Hartmut Grasemann, Christine Bear, Jim Hu

## Abstract

Gene therapy has been the subject of extensive research following the advent of gene-editing technologies. Genetic disorders with difficult-to-target tissues, such as cystic fibrosis (CF), still face many challenges in developing efficacious gene therapy. The potential universal approach of gene replacement involves inserting a functional *CFTR* gene after generating DNA double strand breaks using gene editors such as CRISPR/Cas9. However, this strategy has not achieved clinical significance, as CRISPR/Cas9-mediated integration of *CFTR* is limited primarily by the infrequent activity of the homology-directed repair (HDR) pathway. To circumvent this limitation and improve *CFTR* transgene integration and expression, we explored a method of adding a second integration site, which we termed the dual-locus-targeting method. Using a helper-dependent adenoviral vector (HDAd)-delivered CRISPR/Cas9 system in porcine epithelial cells, we found that sequential delivery of two vectors, one targeting the *CFTR* locus and the other the genomic safe harbour site *GGTA1*, enhanced the integration efficiency of *lacZ* and *CFTR* donor genes to 16.5% and 3.4%, respectively. These results demonstrated a potential strategy to improve the efficacy of *CFTR* replacement for the development of a universal and permanent gene therapy treatment for CF lung disease.

**GRAPHICAL ABSTRACT:** 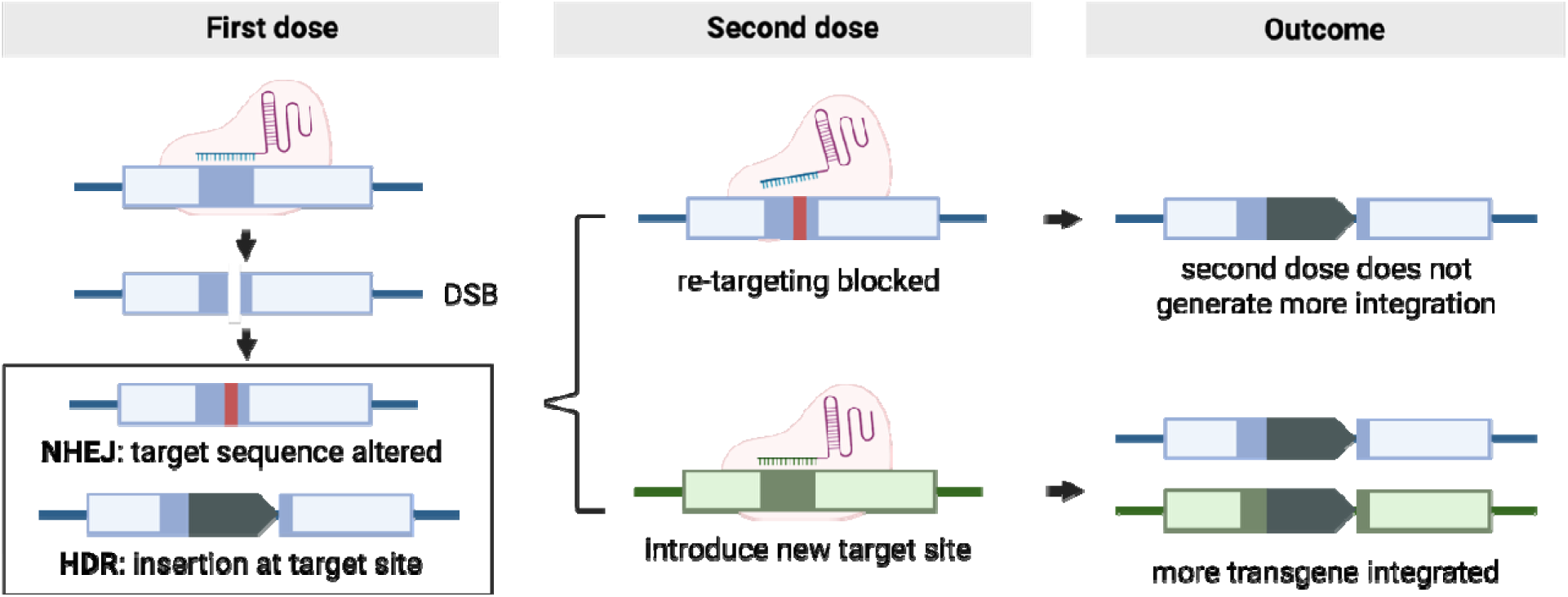

## INTRODUCTION

Cystic fibrosis (CF) is a common inherited disease affecting individuals across all populations. It is caused by mutations in the cystic fibrosis transmembrane conductance regulator (*CFTR*) gene, which encodes an anion channel localized to the apical membrane of epithelial cells in multiple organs [1–3]. CFTR regulates epithelial electrolyte transport; thus, its dysfunction in the airway decreases chloride and bicarbonate export, leading to reduced water efflux, dehydration of the airway surface liquid and subsequent lumenal obstruction. More than 4,000 CFTR variants have been identified, with over 1,000 shown to be disease-causing (cftr2.org). These CF-causing mutations can be categorized into six functional classes based on their effects on CFTR synthesis, processing, and channel activity [4, 5]. Class I mutations introduce premature stop codons that result in truncated mRNAs and, therefore, loss of protein production. The most prevalent CF-causing variant in Caucasian populations, F508del, is a Class II mutation that leads to misfolded CFTR protein, not properly processed or trafficked to the apical cell membrane. Class III to Class VI variants produce CFTR protein but with reduced channel function [4, 5].

CFTR mutations have a major impact on the airways by disrupting mucociliary clearance, which normally helps reduce mucus accumulation and remove inhaled pathogens. As a result, CF airways are susceptible to persistent bacterial infections. Progressive decline in lung function remains the primary cause of morbidity and mortality in CF patients [6]. Most CF patients now experience improved outcomes with CFTR modulators, which are small molecules that target mutant CFTR proteins directly, acting as either correctors that improve folding and trafficking or as potentiators that enhance channel gating. Combination modulator therapies incorporating correctors and a potentiator have been approved for eligible patients [7]. However, because the modulator therapies act at the protein level, their effects are transient and require daily administration. In addition, approximately 10% of patients are ineligible for, not responsive or intolerant to current modulators. Therefore, gene-based therapies are being actively explored as a potentially durable and broadly applicable alternative.

Early gene delivery studies using WT *CFTR* DNA or mRNA demonstrated that exogenous WT *CFTR* nucleic acids can produce functional protein and restore CFTR function in mutant cells and CF animal models [8–10]. However, these non-integrating nucleic acids provide only temporary restoration, as they become diluted during epithelial cell division and turnover, which occur more frequently in CF airways where chronic inflammation accelerates epithelial injury and regeneration. Thus, permanent repair requires two key conditions: (1) integration of the therapeutic cassette into genomic DNA for sustained CFTR expression; and (2) targeting of airway basal stem cells to ensure long-term supply of corrected cells even after turnover, enabling gradual replenishment of the airway with CFTR-expressing cells. CRISPR/Cas9 (Clustered Regularly Interspaced Short Palindromic Repeats and CRISPR-associated protein 9)-based genome editing has emerged as a leading approach to meet the first requirement [11–14].

The CRISPR/Cas9 system consists of a guide RNA (gRNA) that determines the target site and a Cas9 endonuclease that creates a DNA double-strand break (DSB) at the target site [15]. DSBs are repaired via endogenous DNA repair pathways, primarily through the non-homologous end joining (NHEJ) pathway or the homology-directed repair (HDR) pathway. Since HDR only occurs during the S and G2 phases when a sister chromatid is present as a repair template, most DSBs are repaired by the error-prone NHEJ pathway, which often generates indel mutations at the target site [16]. While CRISPR-mediated editing can introduce insertions ranging from single nucleotides to large cDNA cassettes (e.g. ∼4.5 kb CFTR), its efficiency is limited by the low frequency of HDR. Nonetheless, HDR-mediated knock-in (KI) is essential for the universal therapeutic strategies aiming to knock-in a complete CFTR expression cassette capable of benefiting patients regardless of mutation class. Therefore, overcoming the low HDR-dependent integration efficiency remains a central challenge in CF lung gene therapy.

We developed a CRISPR/Cas9-mediated KI system to insert a functional *CFTR* expression cassette into the porcine genome, as the CF pig model closely recapitulates human airway phenotype and pathophysiology (Fig. 1A) [17]. The donor sequence is a previously established *K18-CFTR* cassette, in which human *CFTR* cDNA is driven by a modified human cytokeratin 18 (*KRT18*) promoter [18]. Since *KRT18* is predominantly expressed in epithelial cells, the *K18* promoter enables broad yet tissue-specific expression of CFTR in airway-relevant cell types [19]. *K18-CFTR* has been shown to restore CFTR activity in both *in vitro* and *in vivo* models [8, 9, 20–22]. However, the large size of both CRISPR/Cas9 and *K18-CFTR* cassettes, including homology arms, exceeds the packaging capacity of commonly used gene therapy viral vectors, such as adeno-associated virus (AAV), lentivirus, and retrovirus. Thus, we used a helper-dependent adenoviral (HDAd) vector, which lacks all viral coding sequences, thereby reducing immunogenicity and enabling a large payload capacity of approximately 37 kb. Using this system, our team previously achieved approximately 10% integration efficiency at the *GGTA1* genomic safe harbour locus in porcine cells [22]. However, such *in vitro* efficiencies are often reduced *in vivo* due to biological barriers, including mucus accumulation and immune responses, highlighting the need for further optimization.

**Figure 1.**
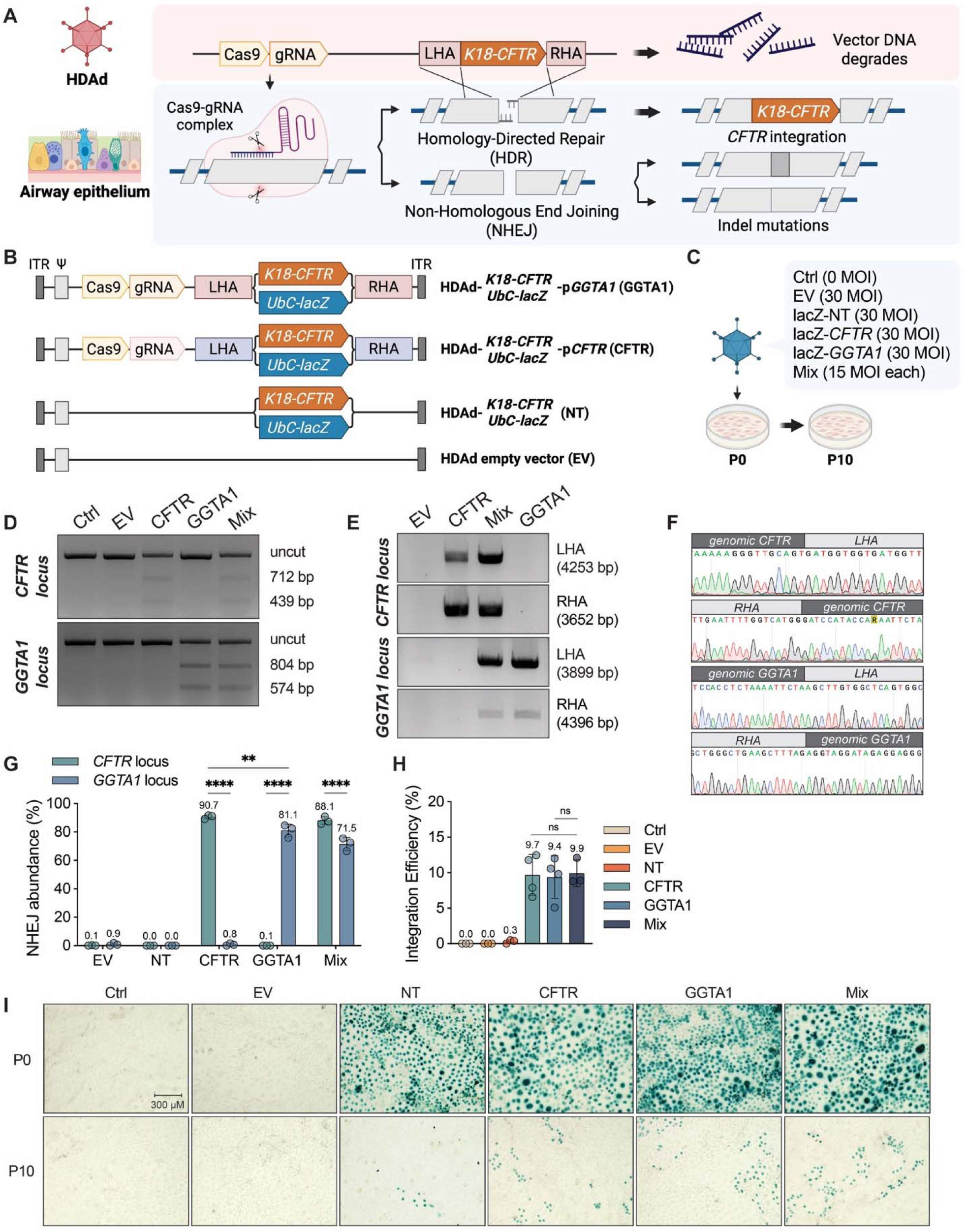
Simultaneous Cotransduction of HDAd-*UbC-lacZ* vectors achieved integration at both CFTR and GGTA1 loci in IPEC-J2 cells. **(A)** Mechanism of CRISPR-mediated integration following HDAd delivery using *K18-CFTR* as an example. DNA double-stranded breaks (DSB) created by Cas9-gRNA complex are repaired by either homology-directed repair (HDR) or non-homologous end joining (NHEJ). If by the HDR pathway, the left and right homology arms (LHA and RHA) allow the *K18-CFTR* donor gene to be inserted at the target site. *K18-CFTR* is an expression cassette with a human CFTR minigene driven by a K18 promoter. Also, vector DNA will likely lose integrity and degrade after recombination. On the other hand, the error-prone NHEJ repair tends to result in indels (insertions and deletions) at the target site. HDAd, helper-dependent adenovirus; sgRNA, single guide RNA. **(B)** HDAd vectors used in this study, two targeting vectors carrying either *UbC-lacZ* or *K18-CFTR* donor gene for each locus (CFTR and GGTA1), non-targeting vectors without the CRISPR/Cas9 gene editing cassette (NT), as well as an empty vector (EV) that only contains inverted terminal repeats (ITR) and Ad5 packaging signal (Ψ) (not shown). The donor genes are flanked by left and right homology arms (LHA and RHA) designed for homologous recombination at each locus during DNA repair. Vectors containing *UbC-lacZ* were used in Fig. 1, 2, and 6, whereas vectors containing *K18-CFTR* were used in Fig. 3, 5, and 7. **(C)** Schematic of experiment design. IPEC-J2 cells were transduced with 30 MOI of HDAd vectors and subcultured to passage 10 (P10) to assess integration. Mix, codelivery of CFTR- and GGTA1-targeting vectors (15 MOI each); gDNA, genomic DNA. **(D)** T7E1 mismatch assay gel showing the expected pattern of fragments after digestion of PCR amplicons from the *CFTR* and *GGTA1* target loci. **(E)** Junction PCR confirming presence of integration at both loci **(F)** Sequencing results of the integration junctions **(G)** frequencies of NHEJ events measured by digital droplet PCR. **(H)** Integration frequencies calculated from X-gal staining images using P10 percentages normalized to P0 percentages. Cell numbers were manually counted from 4 randomly taken images for each group. **(I)** X-gal staining of P0 and P10 cells (10X). *\*\*p < 0.01, ****p < 0.0001, ns = not significant; n = 3. (A-C) were created with BioRender.com*.

A key limitation of CRISPR/Cas9-mediated integration is its reliance on HDR. Most DSBs are repaired via the error-prone NHEJ, resulting in indels that can alter the target sequence and prevent further targeting. We therefore hypothesized that overall integration efficiency could be increased by introducing a second target site. If 10% integration is achievable at a single locus, parallel targeting of an additional locus could theoretically increase total integration up to 20%. Therefore, we investigated whether this dual-locus-targeting strategy enhances integration efficiency. Based on evidence that mutant CFTR can induce unfolded protein response (UPR) and endoplasmic reticulum stress that impacts the production of WT CFTR, we selected the endogenous *CFTR* locus as the second target site [23, 24]. In this strategy, the *K18-CFTR* transgene is inserted into intron 1 of the *CFTR* gene, thereby disrupting expression of mutant CFTR, while simultaneously introducing a functional copy.

We found that sequential delivery of two HDAd vectors significantly improved integration efficiency compared with simultaneous delivery, as measured using the *UbC-lacZ* reporter system. To evaluate functional correction, we generated a W1282X^−/−^ mutant cell line from PK15, a non-CF porcine kidney epithelial cell line, and detected significantly increased CFTR expression using the dual-locus-targeting approach combined with transduction and integration enhancer treatment. These findings suggest that sequential dual-locus-targeting improves integration of both reporter and therapeutic and may represent a generalizable strategy to enhance genome editing efficiency in gene replacement studies.

## MATERIAL AND METHODS

### CRISPR design and plasmid construction

The plasmids for HDAd vectors were cloned into a modified pC4HSU backbone with sequences between NotI and PshAI removed (Fig. 1B) [25]. The donor gene was *lacZ* under *polyubiquitin C (UbC)* promoter or human *CFTR* cDNA under the modified human *KRT18* promoter (*K18*) [18]. The gRNAs were first cloned into pSpCas9(BB)-2A-Puro (Addgene, PX459). The gRNA target sequences were: 5’-GAAGCAAATGACATCACCGC-3’ for *CFTR* locus, and 5’-TTGACATTCATTATTTTCTC-3’ for *GGTA1* locus. The Cas9-gRNA cassettes and homology arms were assembled into the pC4HSU-PN carrying the donor genes.

The plasmids for W1282X^−/−^ cell line generation are pCRISPR-*W1282X*-*EGFP* and pCRISPR-*W1282X*-*mCherry*. Cas9 under *Chicken β-Actin* (*CBA*) promoter, homology arms (approximately 3 kb) and reporter genes (*EGFP* and *mCherry* under *Cytomegalovirus* (*CMV)* promoter, both around 4.7 kb) were cloned into a pTwist-Amp-High-Copy backbone (3 kb) containing the gRNA under *U6* promoter and the *CFTR W1282X* mutant sequences from Twist Bioscience. The gRNA target sequence was: 5’-ATTCAATAACTTTGCAACAG-3’.

### Cell culture

The porcine intestinal epithelial cell line, IPEC-J2, was cultured in DMEM/F12/HAM (Gibco) with 5% porcine serum (Gibco), 1% insulin-transferrin-selenium (ITS-G; Thermo Fisher Scientific), 5 ng/mL epidermal growth factor recombinant human protein (EGF, Thermo Fisher Scientific), and 1% penicillin–streptomycin (10,000 U/mL; Thermo Fisher Scientific). The porcine kidney epithelial cell line, PK15 (WT or W1282X^−/−^), was cultured in MEM (Gibco) with 10% fetal bovine serum (Gibco) and 1% penicillin–streptomycin. All cells were cultured at 37 °C and 5% CO_2_ in a standard tissue culture incubator.

### Transfection

Cells were seeded to obtain a confluence of 70% the next day for transfection (approximate seeding density of 0.38×10^6^ cells). Plasmid DNA was added to the cell culture according to the manufacturer’s instructions (Polyplus jetPRIME, 101000001). The medium was replaced after 4 hours.

### Transduction

Cells were seeded on 6-well plates to obtain a confluence of 50% the next day for transduction (approximate seeding density of 0.27×10^6^ cells). Viral vector was first mixed with 1041 ng of DEAE-dextran per 100 MOI (Multiplicity of Infection) HDAd vectors in 0.8 mL serum-free and antibiotic-free medium per well to enhance transduction efficiency. The transduction mixture was incubated at 37°C for 30 min before adding to the plate. After 4 hours of transduction at 37°C, medium with 10% serum was added to a final volume of 2 mL per well. The vector-containing medium was replaced with fresh growth medium the next day. Small molecules, such as NaBu (Sigma B5887, 2 mM) and M3814 (TargetMol, 3 μM), were added at seeding and removed at day 2 post-transduction to enhance integration. Other molecules tested include Trichostatin A (Sigma T8552), SCR7 (STEMCELL 74102), RS-1 (STEMCELL 74092), and L755507 (STEMCELL 73992).

### Fluorescence-Activated Cell Sorting (FACS)

Transfected cells were trypsinized and resuspended in stain buffer supplemented with FBS (BD Pharmingen, 554656). Cells were stained with viability dye (Live/Dead Fixable Violet Dead Cell Stain Kit, Thermo Fisher Scientific) to gate out the dead cell population during sorting, followed by filtering through nylon mesh. The single cell suspension was sorted into 96-well plate with 70 μl of conditioned medium (MEM collected from PK15 cell culture, filtered, and supplemented with FBS to a final percentage of 20%) at 1 cell per well using a Sony MA900 VBYR sorter at the SickKids-UHN Flow Cytometry Core Facility. Sorted cells were recovered and cultured in regular MEM medium with 10% FBS.

### Membrane Potential Assay (FLIPR)

Cells were cultured on a 96-well black clear-bottom plate (Costar 3603) to reach 100% confluency. After 30 min incubation at 37°C in 0.5 mg/ml membrane potential dye (FLIPR Membrane Potential Assay Kit, Molecular Devices) prepared in NMDG-gluconate buffer (150 mM NMDG-Gluconate, 3 mM potassium gluconate, 150 mM gluconate lactone, 10 mM HEPES, pH 7.35, osmolarity 300 mOsm). Fluorescence measurements were taken at 37°C (excitation: 530 nm, emission: 560 nm) with a SpectraMax i3X Multi-mode Assay Microplate Reader (SickKids Hospital Structural and Biophysical Core Facility) at 30s intervals, with 11 reads for baseline, optional 11 reads of amiloride (50 μM in DMSO, ThermoFisher, J62168.03) to block sodium channel activity, 21 reads after forskolin (FSK, 10 μM in DMSO, Sigma-Aldrich, F3917) stimulation, and 15 reads after CFTR_inh_-172 (inh172, 10 μM in DMSO, Sigma-Aldrich, C2992) inhibition. DMSO was used as a vehicle control for forskolin. IBMX (200 μM in DMSO, Sigma-Aldrich, I5879) was occasionally added together with forskolin to boost channel activity.

### Real-Time Quantitative PCR (RT-qPCR)

RNA samples were collected at day 2 post-transduction and extracted using Amersham RNAspin Mini Kit (Cytiva 25050072). One μg of total RNA from each sample was reverse-transcribed using random hexamers by SuperScript IV VILO Master Mix kit (ThermoFisher Scientific, 11755050) following the manufacturer’s protocol. Ten μg of cDNA were used as the template for qRT-PCR using SYBR Green PCR Master Mix (Applied Biosystems, A25742) on a QuantStudio 3 Real-Time PCR System (Applied Biosystems). Primers used are listed in Table S1. Fold changes were calculated with the comparative Ct method.

### Junction PCR and sequencing

Genomic DNA was isolated using the QIAamp DNA mini kit (QIAGEN, 51306). The primers were designed to flank the homology arm junctions, so that one primer binds to sequences unique in genomic DNA and the other primer binds to sequences unique in vector DNA. The same primers were used as listed in Table S1.

### T7E1 Mismatch Assay

Genomic DNA was isolated using the QIAamp DNA mini kit (QIAGEN, 51306). DNA was amplified using these primers. The PCR products were denatured and reannealed, followed by T7E1 digestion (NEB M0302) at 37°C for 30 min. The fragments were resolved on a gel. The PCR primers were used as listed in Table S1. The expected fragments are 765 bp and 243 bp for the *CFTR* locus, 574 bp and 804 bp for the *GGTA1* locus.

### Digital Droplet PCR (ddPCR)

Digital droplet PCR was performed as previously described at the Centre for Applied Genomics of the Hospital for Sick Children [22]. Genomic DNA was first PCR-amplified using the forward and reverse primers in Table S1. To measure indel mutations, two probes were designed to detect sequences within the amplicons: a VIC-labeled probe (green-yellow fluorescent dye) binding the unedited target sequence, and a FAM-labeled probe (bright green fluorescent dye) binding a region outside the target site, which serves as a reference for total amplicon copy number. Indel mutations at the target site resulting from NHEJ repair prevent VIC probe binding, enabling quantification of NHEJ-derived alleles.

To measure integration, genomic DNA was isolated from cells at Passage 10 and amplified with the forward and reverse primers in Table S1. Two VIC probes were designed, one binding *K18-CFTR* and the other binding the HDAd packaging signal (Ad5). The FAM probe detects a porcine reference gene *Tfrc*. The two VIC probes are used in separate reactions with the FAM probe to detect the abundance of K18-CFTR and the remaining viral genome in the cells. Since the residual viral genome may carry unintegrated K18-CFTR, the integration efficiency of K18-CFTR is calculated by subtracting the viral genome copy number.

### Western Blot

Proteins were isolated by cell lysis with RIPA buffer (1% Triton X-100, 0.1% SDS, 150 mM NaCl, 20 mM Tris-HCl, 0.5% Deoxycholate) with 10% protease inhibitor. Protein concentrations were measured by BCA assay (Pierce bicinchoninic acid assay [BCA] Protein Assay kit; ThermoFisher Scientific). BCA assay was quantified with the VarioSkan LUX Plate reader from the Imaging Facility at the Hospital for Sick Children. Cell lysates were mixed with 4× Laemmli sample buffer (Bio-Rad, 1610747, final concentration of 1×) and β-mercaptoethanol (Sigma-Aldrich, final concentration of 2.5%). Protein samples (30-50 μg) were loaded onto mini-PROTEAN TGX Stain-Free Gels (Bio-Rad, #4568086). For CFTR protein detection specifically, cell lysates were mixed with 4× sample buffer to a final concentration of 2× and with β-mercaptoethanol to a final concentration of 5%, then loaded directly onto the gel without incubation at 95°C. Protein bands were transferred onto Amersham Protran Premium 0.45 μm nitrocellulose membrane (GE Healthcare Life Science, 10600003), which was then probed with anti-CFTR antibody (596, 1:1000 dilution), or anti-GAPDH antibody (Novus Biologicals, NB300-328, 1:5000 dilution), followed by goat anti-mouse IgG (H+L)-HRP conjugate antibody at 1:2000 and 1:5000, respectively (Bio-Rad, 170-6516). Antibodies were prepared in a blocking solution of 5% nonfat dry milk (Bio-Rad, 1706404) in TBST. TBST was diluted from 10× Tris Buffered Saline (Multicell, 02962) and supplemented with 0.5 M Tween-20 (Fisher, BP337-500).

### HDAd vector production

HDAd vectors were produced as previously described [26, 27]. Vectors were amplified by serial coinfections of producer cells (116) with the HDAd and the helper virus (NG163) from one 6 cm plate to one 10 cm plate, to one 15 cm plate, to 8×15 cm plates, and finally to a 3 L flask. HDAd was rescued from the 3 L infected cell culture. After cell lysis, HDAd was purified by CsCl density gradient centrifugation. The viral titer was measured by absorbance at 260 nm as well as real-time quantitative-PCR.

### X-gal staining

X-gal staining was performed at day 2 post-transduction. Cells on 6-well plates were fixed with 5% glutaraldehyde for 15 min at room temperature and stained with staining solution (0.8mg/ml X-gal, 2 mM MgCl_2_, 5 mM potassium ferricyanide, and 5 mM potassium ferrocyanide) for 3 hours or overnight at 37°C, depending on the *lacZ* expression level. Solutions were prepared in washing buffer (sodium phosphate buffer, pH 7.8). Cells were washed 3 times before and after each step. Random images were taken for each well after staining. The number of blue cells (lacZ-expressing cells) was counted manually from four images per well.

### Off-target analyses

CHOPCHOP was used to predict potential off-target sites [28]. Out of the 46 sites with a distance of 3 from the target sequence, we selected 4 sites for analysis. Genomic DNA was isolated from PK15 W1282X^−/−^ cells that received the CFTR+GGTA1 vector. Primers were designed to generate an approximately 400-500 bp amplicon surrounding the off-target sites (Table S1). One of the primer pair was used for sequencing, and the results were analyzed using the Inference of CRISPR Edits (ICE) tool for the frequency of edits [29].

### Statistical analyses

Statistical significance was calculated by one-sample t-test or one-way/two-way ANOVA using GraphPad Prism 9 software. A p-value <0.05 was considered statistically significant. Tukey’s test was used for multiple comparisons after ANOVA. Data and error bars represent mean ± standard deviation of n = 3-6 independent biological samples for each cell line.

## RESULTS

### Simultaneous delivery of CFTR- and GGTA1-targeting vectors in IPEC-J2 cells

To build on the previously reported GGTA1-targeting vector and determine whether the addition of a CFTR-targeting vector could increase HDR-mediated integration, we generated two CFTR-targeting HDAd vectors carrying either the *UbC-lacZ* reporter or the *K18-CFTR* donor cassette (Fig. 1A, B). *UbC-lacZ* vectors were first used to establish proof of concept, followed by *K18-CFTR* vectors to assess functional restoration. For simplicity, vectors are hereafter referred to by their target loci as “CFTR” and “GGTA1” vectors (Fig. 1B). To examine whether co-transduction of these two vectors improves integration efficiency, non-CF porcine intestinal epithelial cells IPEC-J2 were transduced with a total of 30 MOI (Multiplicity of Infection) of HDAd vectors (Fig. 1C). An empty vector (EV) and a non-targeting lacZ vector lacking Cas9 (NT) were included as controls (Fig. 1C). To confirm the presence of DNA DSB repair products generated through NHEJ and HDR, we performed the T7 Endonuclease 1 (T7E1) mismatch assay and junction PCR to detect indel mutations and targeted integrations, respectively. Genomic DNA isolated from transduced cells was amplified across the target site. As expected, T7E1-generated small fragments were detected at both *CFTR* and *GGTA1* loci in cells transduced with the corresponding vectors, confirming successful Cas9 cleavage (Fig. 1D). Junction PCR further demonstrated *lacZ* integration at both loci, and amplicon sequencing confirmed insertion at the expected sites (Fig. 1E, F).

To quantify editing outcomes, ddPCR was used to measure indel frequencies and X-gal staining was used to estimate integration efficiency (Fig. 1G-I). Indel frequencies reached approximately 90% at the *CFTR* locus, and over 70% at the *GGTA1* locus (Fig. 1G). Since both T7E1 mismatch assay and ddPCR interrogate only a small genomic region, they do not detect alleles with large insertions. Nevertheless, the results indicate that at least 70% of amplifiable alleles underwent NHEJ repair, while fewer than 30% remained unedited. Although rare, some unedited alleles detected may have undergone precise NHEJ repair [30]. Together, these data indicate highly efficient Cas9 cleavage at both loci. X-gal staining was then used to visualize lacZ-expressing cells at passage 0 (P0) and passage 10 (P10) (Fig. 1H, I). We assumed that most unintegrated *lacZ* copies were diluted to negligible levels by P10 and that the remaining lacZ-positive cells represent stable integrations. As expected, integration efficiencies in the EV and NT controls were minimal, whereas Cas9-containing vectors produced substantially higher levels of integration (Fig. 1H). Although co-transduction of CFTR and GGTA1 vectors yielded the highest integration of 9.9% (Fig. 1H, “Mix”), this was not significantly different from the single-locus groups (9.7% for “CFTR” and 9.4% for “GGTA1”). Collectively, these results demonstrate that the CFTR-targeting vector exhibited efficient Cas9 cleavage efficiency and an integration efficiency comparable to that of the GGTA1-targeting vector, but simultaneous transduction of both vectors did not significantly improve *lacZ* integration efficiency.

### *UbC-lacZ* integration efficiency can be increased by sequential delivery of CFTR- and GGTA1-targeting vectors

Since HDR occurs only during the S and G2 phases of the cell cycle, we hypothesized that co-transduction or simply increasing the viral doses would not further improve the integration efficiency because the number of cells capable of HDR is limited. Assuming that approximately 10% of cells are in S/G2 phase at the time of transduction, delivering additional vectors cannot increase integration beyond this proportion. In contrast, if HDAd vectors are delivered at separate time points, it is unlikely that the same subset of cells will be in S/G2 phase during both transductions. Sequential delivery may therefore increase the total number of cells available for HDR-mediated integration. To test this hypothesis, we evaluated sequential transduction in both IPEC-J2 and PK15 (non-CF porcine kidney epithelial) cells.

Cells were first transduced with 30 MOI of lacZ-expressing vectors at P0 (Fig. 2A, “NT”, “CFTR”, and “GGTA1”), followed by a second 30 MOI dose at P3 (Fig. 2A). To compare dual-locus-targeting versus repeated single-locus targeting, cells receiving CFTR or GGTA1 vector at P0 were divided into three groups and transduced at P3 with NT, CFTR, or GGTA1 vector (Fig. 2A). Cells were subsequently expanded to P10, and stable lacZ integration was assessed by X-gal staining (Fig. 2B, C). The percentages of lacZ-positive cells at P0 and P10 were quantified manually (Fig. 2D-G). In IPEC-J2 cells, transduction with the CFTR vector at P0 followed by the NT vector at P3 resulted in an integration efficiency of 6.8% (Fig. 2E, CFTR+NT). As expected, a second dose of CFTR vector (CFTR+CFTR) did not significantly increase integration. In contrast, cells receiving CFTR followed by GGTA1 vector (CFTR+GGTA1) achieved a significantly higher integration efficiency of 16.5% (Fig. 2E). Similar results were obtained when the order of HDAd delivery was reversed (GGTA1+CFTR), indicating that the order of transduction did not affect integration efficiency and supporting the sequential dual-locus-targeting (Fig. 2E, GGTA1+CFTR).

**Figure 2.**
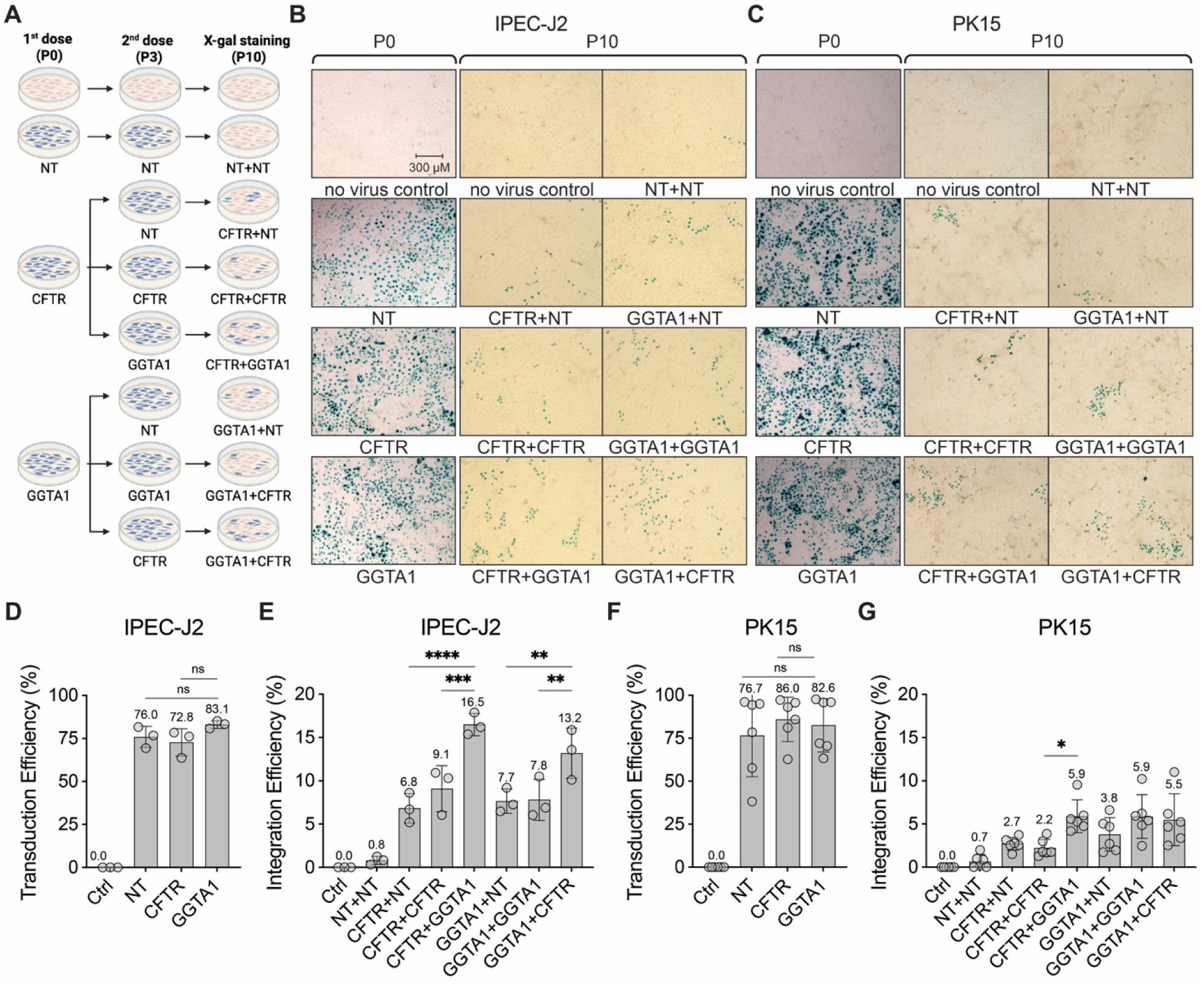
Sequential delivery of the CFTR- and GGTA1-targeting HDAd-*UbC-lacZ* vectors in IPEC-J2 and PK15 cells improves integration efficiency. **(A)** Schematic of sequential HDAd delivery. IPEC-J2 and PK15 cells received 30 MOI of HDAd vector at both P0 and P3. “CFTR+GGTA1” represents the first dose targeting the *CFTR* locus and the second dose targeting the *GGTA1* locus. *Created with BioRender.com.* (B, C) Images of X-gal staining at P0 and P10 from IPEC-J2 cells (B) and PK15 cells (C). **(D-G)** Transduction and integration efficiencies were calculated from the above images for IPEC-J2 cells (D, E) and PK15 cells (F, G). Ctrl, no virus control; EV, empty vector; NT, no target (no Cas9 control); MOI, multiplicity of infection. *Ordinary one-way ANOVA, *p < 0.05, **p < 0.01, ***p < 0.001, ****p < 0.0001, ns = not significant; n = 3 for IPEC-J2, n = 6 for PK15*.

Overall integration efficiencies were lower in PK15 cells than in IPEC-J2 cells, consistent with previous observations and likely reflecting cell-type-specific differences (Fig. 2G). Therefore, six biologically independent replicates were performed. Among the treatment groups, only CFTR+GGTA1 showed significantly higher integration efficiency than CFTR+NT and CFTR+CFTR (Fig. 2G). In addition, the GGTA1 vector appeared to integrate slightly more efficiently than the CFTR vector in PK15 cells, as cells receiving the GGTA1 vector generally exhibited higher integration rates. Nevertheless, addition of the CFTR vector further increased the overall integration efficiency (Fig. 2G, CFTR+GGTA1 versus GGTA1+NT). Together, these findings indicate that CRISPR/Cas9-mediated integration is constrained by the limited number of cells capable of HDR-mediated repair. However, integration efficiency can be improved through sequential delivery of the dual-locus-targeting vectors, which increases the number of cells that can undergo HDR.

### Dual-locus-targeting strategy enhances the integration of *K18-CFTR* in IPEC-J2 cells

After establishing proof of concept with *UbC-lacZ*, we next evaluated the dual-locus-targeting strategy using the therapeutic *K18-CFTR* donor cassette (Fig. 1B). Since the CFTR+GGTA1 combination yielded the highest integration efficiency with *UbC-lacZ* in IPEC-J2 cells, we used the same delivery order for *K18-CFTR*. Although IPEC-J2 is a non-CF cell line, its low endogenous CFTR expression may permit detection of transgene-derived CFTR expression above baseline levels following successful *K18-CFTR* integration. Experimental controls included a no-transduction control (Ctrl), an empty vector control (EV+EV), a non-targeting *K18-CFTR* control (NT+NT), and single-locus-targeting controls (CFTR+NT and GGTA1+NT) (Fig. 3A).

**Figure 3.**
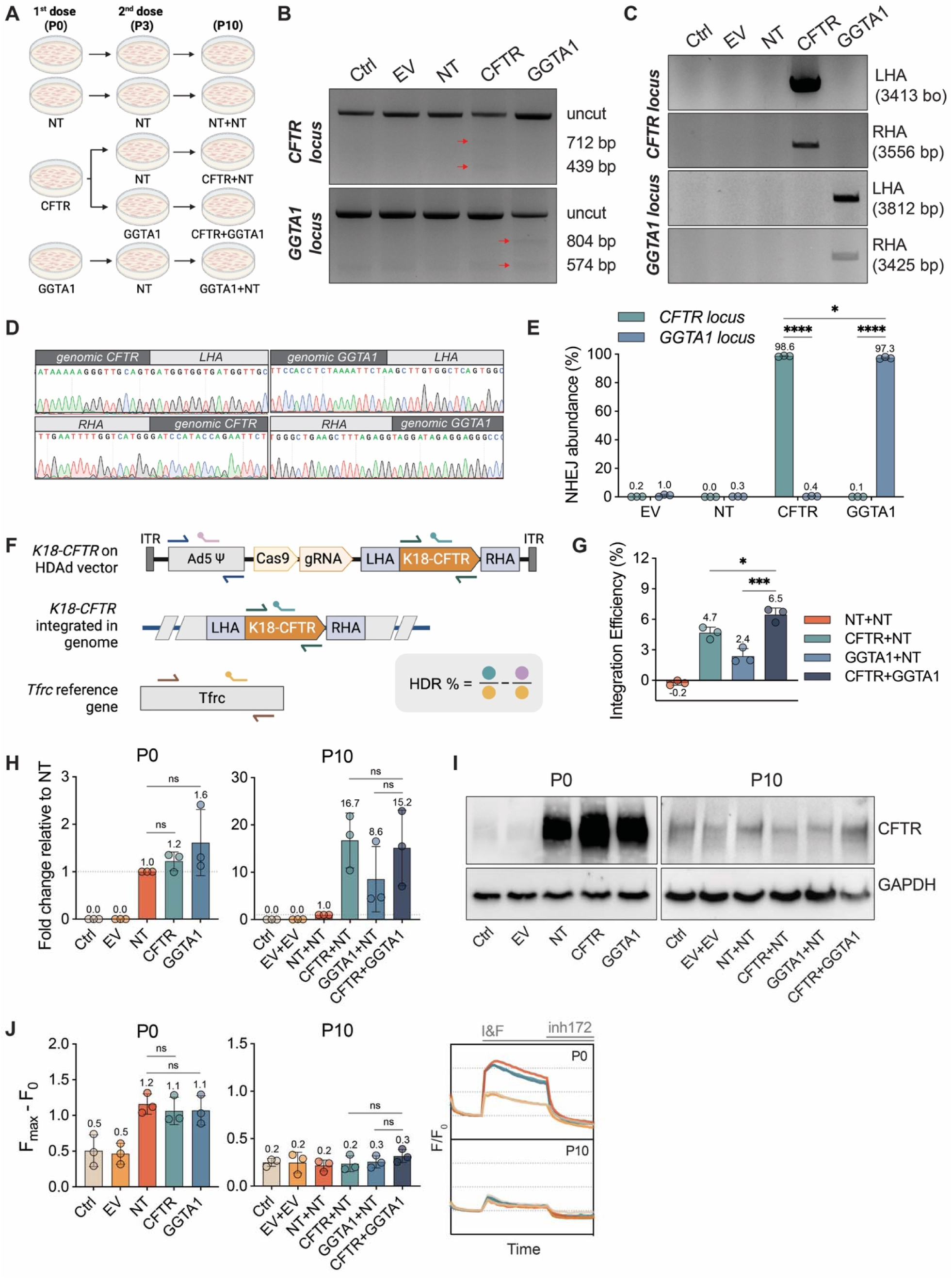
CFTR integration and expression in IPEC-J2 cells following dual-locus-targeting HDAd delivery. **(A)** Schematic of experiment timeline. **(B)** T7E1 mismatch assay gel showing the expected pattern of fragments (indicated by the red arrows) after digestion at both *CFTR* and *GGTA1* loci. **(C)** Junction PCR analyses confirming the presence of the integrated *CFTR* donor gene at both target sites. **(D)** Sequencing of the junction PCR products at each junction. **(E)** NHEJ abundance measured using ddPCR. **(F)** Schematic of probe design for ddPCR to detect integration. **(G)** Integration efficiency measured with ddPCR. **(H)** Transcript level of *K18-driven CFTR* relative to “NT”, measured with RT-qPCR at P0 and P10. **(I)** Western blot of CFTR protein at P0 and P10 with GAPDH as the loading control. **(J)** CFTR function measured by FLIPR assay at P0 and P10*. Left two panels:* the highest increase in fluorescence (F_max_-F_0_) after forskolin (FSK) activation. *Right panel:* change in baseline-normalized membrane fluorescence (F/F_0_) recorded over 27 min with IBMX and FSK (I&F) added at 5 min, and CFTRinh-172 (inh172) added at 15 min. Ctrl, untransduced control; EV, empty vector; NT, non-targeting vector with K18-CFTR but not Cas9; ITR, inverted terminal repeats; Ψ, Ad5 packaging signal. *Two-way ANOVA*; *\*p < 0.05, ***p < 0.001, ****p < 0.0001, ns = not significant; n = 3*. *(A) and (F) were created with BioRender.com*.

To determine the optimal viral dose, *K18-CFTR* vectors were first evaluated in a dose-response test measuring CFTR channel activity by membrane potential assay (FLIPR, FLuorescence Imaging Plate Reader; data not shown). Based on these results, IPEC-J2 cells were transduced with 100 MOI of HDAd at both P0 and P3, and CFTR expression was assessed at P10. To confirm the generation of NHEJ and HDR-mediated repair products, T7E1 mismatch assay and junction PCR were performed using genomic DNA collected at P0. T7E1 analysis demonstrated efficient Cas9 cleavage at both *CFTR* and *GGTA1* loci, while junction PCR confirmed *K18-CFTR* integration at the corresponding target sites (Fig. 3B, C). Amplicon sequencing further verified the correct insertion of *K18-CFTR* at the intended genomic locations (Fig. 3D). Following validation of vector functionality, ddPCR was used to quantify NHEJ and HDR frequencies. Indel frequencies reached 98.6% and 97.3% at the *CFTR* and *GGTA1* locus, respectively, indicating highly efficient Cas9 cleavage (Fig. 3E). These values are consistent with those observed using the *UbC-lacZ* vectors, as both systems employ the same Cas9-gRNA cassettes (Fig. 1B). Since *K18-CFTR* integration cannot be directly visualized as with *lacZ* reporter, integration efficiency was quantified by ddPCR using a probe specific for the human *CFTR* cDNA sequence present in *K18-CFTR* but absent from endogenous porcine *CFTR* (Fig. 3F). A second probe targeting the HDAd genome was used to quantify the residual episomal donor copies. Both probes were analyzed in separate reactions together with a genomic reference probe (Fig. 3F). *K18-CFTR* integration efficiency was then estimated by subtracting the HDAd genome copies from total *K18-CFTR* copies. Using this approach, single-locus-targeting achieved integration efficiencies of 4.7% and 2.4% at the *CFTR* and *GGTA1* locus, respectively (Fig. 3G). Consistent with the *UbC-lacZ* results, dual-locus-targeting (CFTR+GGTA1) produced the highest integration efficiency, reaching 6.5% (Fig. 3G). These results demonstrated that the dual-locus-targeting strategy also enhanced the integration of the therapeutic K18-*CFTR* transgene.

Next, we examined whether this 6.5% integration resulted in detectable increases in CFTR expression. CFTR expression was assessed at the mRNA, protein, and functional levels in P0 and P10 cells using RT-qPCR (Fig. 3H), Western blotting (Fig. 3I), and FLIPR assay (Fig. 3J), respectively. At P0, CFTR expression primarily originated from *K18-CFTR* carried by the HDAd vectors, whereas expression detected at P10 was expected to reflect stable genomic expression. To distinguish the integrated *K18-CFTR* from the endogenous porcine *CFTR*, RT-qPCR primers were designed to specifically detect the human *CFTR* donor sequence. *K18-CFTR* transcript levels were shown as fold change relative to the non-targeting control (“NT”). As expected, cells receiving NT, CFTR, and GGTA1 vectors showed similar *K18-CFTR* transcript levels at P0 because all groups received equivalent doses of the donor cassette (Fig. 3H). At P10, all groups receiving Cas9-containing vectors exhibited substantially higher fold changes due to the minimal transcript level of “NT”, which comes from remaining NT vectors in the cells or from spontaneous integration of *K18-CFTR* in the absence of Cas9. Thus, these results indicate successful Cas9-mediated integration (Fig. 3H). CFTR protein expression was also assessed by Western blotting. Consistent with the transcript data, CFTR protein was readily detected at P0 in groups receiving K18-CFTR donor but not the no-CFTR controls (Fig. 3I, “Ctrl” and “EV”). At P10, CFTR protein levels were similar among all groups, although the CFTR+GGTA1 consistently showed slightly higher band intensity than the single-locus-targeting groups (Fig. 3I).

CFTR channel activity was assessed using the FLIPR membrane potential assay. At P0, groups receiving K18-CFTR exhibited higher CFTR activity than the endogenous baseline level (Fig. 3J, P0). However, by P10, CFTR function was not significantly different from the endogenous level and did not differ significantly among treatment groups (Fig. 3J, P10). Although the CFTR+GGTA1 group showed a trend toward higher CFTR protein expression and forskolin (FSK)-stimulated channel activity than the single-locus-targeting groups (Fig. 3I, J), these differences were modest. The absence of a substantial increase in CFTR protein expression or function despite a 6.5% integration frequency may be partially due to disruption of endogenous WT *CFTR* alleles following integrating at the *CFTR* locus, or to regulation of genomic CFTR expression at the endogenous levels in this jejunum epithelial cell type (Fig. 3F). Hence, these findings indicate that evaluation of functional CFTR rescue will require a CFTR-deficient model in which endogenous WT CFTR expression does not mask transgene-derived activity.

### Generation of PK15 W1282X^−/−^ cell line for the assessment of CFTR integration

We initially planned to mutagenize a commercially acquired WT porcine tracheal epithelial cell line (PTE) for this study. Unfortunately, functional analysis revealed undetectable CFTR activity in PTE cells, making them unsuitable as a positive control for CFTR restoration studies (Fig. S1A). In addition, HDAd transduction efficiency was low in PTE cells (Fig. S1B). Although IPEC-J2 cells exhibited higher integration efficiencies than PK15 cells in the lacZ experiments (Fig. 2.5E, G), they displayed signs of senescence after extended culture and were therefore considered unsuitable for the repeated single-cell sorting required for mutant cell line generation. Consequently, PK15 cells, which exhibited the highest endogenous CFTR function among the cell lines tested, were selected for subsequent studies (Fig. S1A).

The W1282X mutation is a nonsense mutation that converts TGG to TGA at exon 23 of *CFTR* (Fig. 4A). This mutation was selected because it is located distal to the intron 1 target site used for subsequent *K18-CFTR* integration, minimizing the likelihood that the integration target sequence would be altered in the gene editing process. In addition, Class I mutations such as W1282X produce little or no functional CFTR protein and are therefore largely unresponsive to CFTR modulators, as the absence of functional CFTR protein renders them unresponsive to CFTR modulators. Restoration of CFTR function in a W1282X^−/−^ mutant cell line would therefore provide a relevant proof of concept for this gene replacement strategy. To generate a homozygous W1282X^−/−^ mutation, we designed two CRISPR/Cas9 plasmids containing a repair template with the TGA sequence, homology arms, and either an mCherry or EGFP reporter gene (Fig. 4B). Cells expressing both reporters following editing were expected to carry the mutation in both alleles. Since the WT TGG can serve as a PAM sequence, the gRNA target sequence was designed to be immediately upstream of TGG, enabling Cas9 cleavage three nucleotides before TGG (Fig. 4C). HDR-mediated repair using the donor templates introduced the TGA mutation together with either mCherry or EGFP cassette into the downstream intronic region (Fig. 4C). To prevent interference with future FLIPR analysis, loxP sites were incorporated flanking both reporter genes, allowing their subsequent removal by Cre recombinase while retaining the W1282X mutation and a single residual loxP site (Fig. 4C). The overall workflow consisted of two rounds of single-cell sorting the clone selection, with functional screening by FLIPR assay at each stage (Fig. 4D).

**Figure 4.**
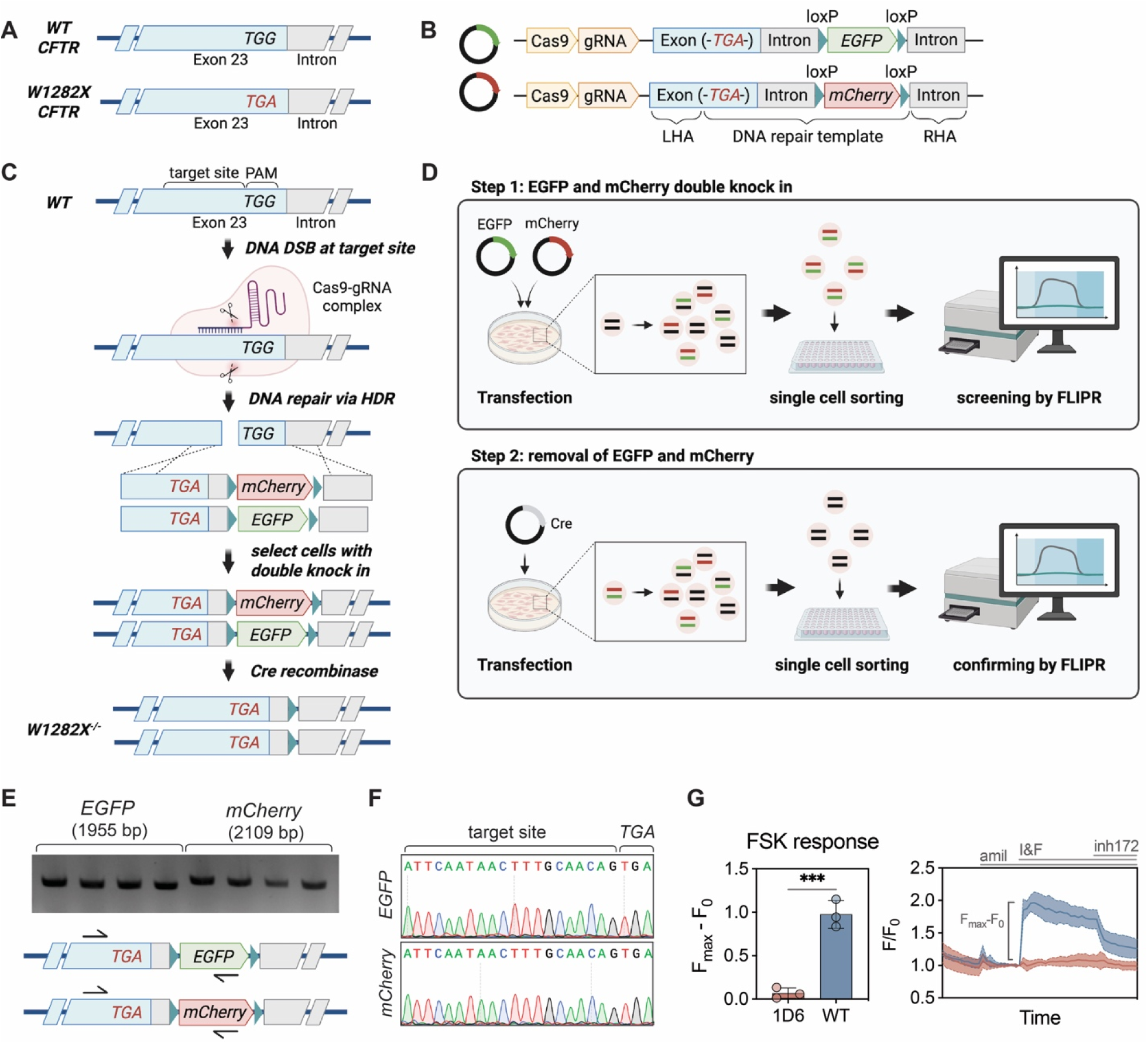
PK15 W1282X^−/−^ cell line generation. **(A)** W1282X is a TGG to TGA nonsense mutation in exon 23 of *CFTR*. **(B)** Plasmid DNAs designed for mutagenesis. mCherry and EGFP were reporters for the presence of the donor containing TGA. The reporter genes are flanked by loxP sites for removal after TGA mutation is achieved. **(C)** Schematics of the mutagenesis process after delivery of the plasmid DNAs into PK15 cells. To create homozygous W1282X mutations, cells expressing both mCherry and EGFP were selected. These reporter genes were later removed with Cre recombinase. **(D)** Schematics of the two selection steps to achieve W1282X double knock-in and the removal of the reporter genes by single-cell sorting. **(E)** Junction PCR with one primer binding the reporter gene and the other binding the genomic DNA, confirming the presence of donor sequences at the target sites**. (F)** Sequencing of the junctions also confirms the correct “*TGA*” edit. **(G)** CFTR function measured by FLIPR assay showing knocked out function in clone 1D6 after the second selection compared to WT PK15 cells. *Left panel:* highest increase in fluorescence (F_max_-F_0_) after forskolin (FSK) activation. *Right panel:* change in baseline-normalized membrane fluorescence (F/F_0_) recorded over 27 min with amiloride (amil) added at 5 min, IBMX and FSK (I&F) added at 10 min, and CFTRinh-172 (inh172) added at 20 min. *One sample t-test*; \*\**\*p < 0.001; n = 3*. *(A-E) were created with BioRender.com*.

PK15 cells were co-transfected with both plasmids, and mCherry/EGFP double-positive cells were isolated by single-cell sorting (Fig. S1C, D). Candidate clones exhibiting low CFTR activity by FLIPR assay were further analyzed by junction PCR (data not shown). PCR primers were designed with one primer upstream of the TGA stop codon and the other within the reporter cassette, such that successful amplification targeted insertion at the corresponding allele (Fig. 4E). All four candidate clones generated amplicons at both mCherry and EGFP junctions, consistent with homozygous mutation, which is also confirmed by sequencing of the junctions (Fig. 4F). Following Cre-mediated reporter removal and a second round of selection, Clone #1D6 was chosen for subsequent experiments. This clone exhibited minimal CFTR function (Fig. 4G), contained the W1282X mutation at exon 23, and retained an intact intron 1 target sequence for *K18-CFTR* integration (Fig. S1E). In conclusion, these results confirmed that Clone #1D6 carries the W1282X^−/−^ mutation and is suitable for evaluating CFTR restoration by the dual-locus-targeting strategy.

### The dual-locus-targeting approach generates low levels of CFTR targeting in PK15 W1282X^−/−^ Cells

As in IPEC-J2 cells, *K18-CFTR*-expressing vectors were first evaluated in PK15 W1282X^−/−^ cells at various doses to determine the MOI required to restore WT-level CFTR function in the FLIPR assay. Similarly, 100 MOI of HDAd vectors were delivered to PK15 W1282X^−/−^ cells at both P0 and P3. Cells receiving the CFTR vector were divided into two groups for CFTR+NT and CFTR+GGTA1 combinations at P3 (Fig. 3A). The functionality of the vectors in PK15 W1282X^−/−^ cells was first confirmed by T7E1 mismatch assay and junction PCR. Indel mutations and K18-CFTR integrations were detected at both target loci, indicating successful Cas9 cleavage and HDR-mediated integration (Fig. 5A, B). Sequencing of the junction PCR amplicons further confirmed integration at the intended genomic locations (Fig. 5C). Next, the DNA repair outcomes were quantified using ddPCR (Fig. 5D, E). Indel frequencies reached 89.2% at the *CFTR* locus and 87.3% at the *GGTA1* locus, indicating efficient Cas9 cleavage with over 10% of amplifiable alleles remained unedited (Fig. 5D). These NHEJ efficiencies were approximately 10% lower than those observed in IPEC-J2 cells (Fig. 3E). Integration efficiencies were below 2% in all groups receiving integrating vectors (Fig. 5E). Changes in CFTR expression between P0 and P10 was assessed by RT-qPCR (Fig. 5F), Western blotting (Fig. 5G), and FLIPR assay (Fig. 5H). Because PK15 W1282X^−/−^ cells lack functional CFTR, CFTR expression detected at P10 is expected to originate primarily from integrated *K18-CFTR*. Consistent with the low integration efficiencies measured by ddPCR, *K18-CFTR* transcript levels at P10 were not significantly higher than those in the non-targeting control (Fig. 5F). Likewise, CFTR protein and channel function remained undetectable in all experimental groups at P10 (Fig. 5G, H). Together, these results suggest that the dual-locus-targeting strategy did not produce a significant increase in *K18-CFTR* integration or detectable CFTR restoration in PK15 W1282X^−/−^ cells. Therefore, additional approaches to enhance integration efficiency are needed to achieve functional CFTR rescue.

**Figure 5.**
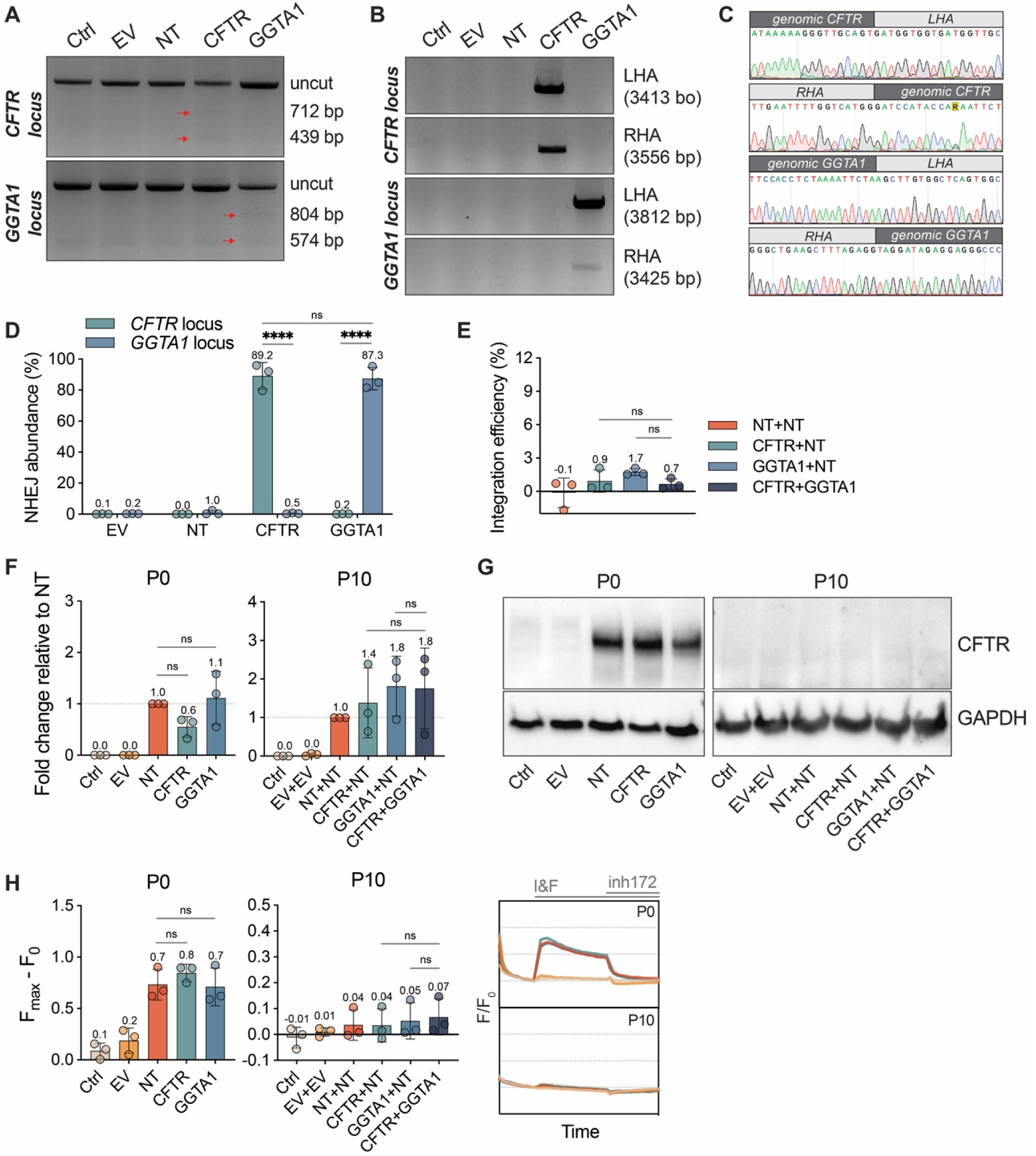
CFTR integration in PK15 W1282X^−/−^ cells following dual-locus-targeting HDAd delivery. **(A)** T7E1 mismatch assay gel showing the expected pattern of fragments (indicated by the red arrows) after digestion at both *CFTR* and *GGTA1* loci. **(B)** Junction PCR analyses confirming the presence of the integrated *CFTR* donor gene at both loci. **(C)** Sequencing of the junction PCR products at each junction. **(D)** NHEJ abundance measured with ddPCR. **(E)** Integration efficiency measured with ddPCR. **(F)** *K18-CFTR* transcript level normalized to “NT”, measured by RT-qPCR at P0 and P10. **(G)** Western blot of CFTR protein at P0 and P10 with GAPDH as the loading control. **(H)** CFTR function measured by FLIPR assay at P0 and P10. *Left two panels:* the highest increase in fluorescence (F_max_-F_0_) after forskolin (FSK) activation. *Right panel:* change in baseline-normalized membrane fluorescence (F/F_0_) recorded over 27 min with IBMX and FSK (I&F) added at 5 min, and CFTRinh-172 (inh172) added at 15 min. Ctrl, untransduced control; EV, empty vector; NT, nontargeting vector with K18-CFTR but not Cas9; LHA and RHA, left and right homology arms. *Two-way ANOVA*; *\*\*\*\*p < 0.0001, ns = not significant; n = 3*.

### Integration efficiencies can be enhanced with NaBu+M3814 treatment in PK15 W1282X^−/−^ cells

To further improve the integration efficiency in PK15 W1282X^−/−^ cells, we evaluated seven small molecules previously shown to enhance HDR-mediated gene integration. L755507 [31, 32] and RS-1 [33, 34] have been identified as HDR-promoting compounds, whereas M3814 [35–37] and SCR7 [38, 39] inhibit the competing NHEJ pathway. We also included the histone deacetylase inhibitors (HDACi) sodium butyrate (NaBu) [40] and trichostatin A (TSA) [41, 42], which increase chromatin acetylation and may enhance genomic accessibility to DNA repair machinery.

To assess the integration efficiency, we employed the easily detectable *lacZ* vector. PK15 W1282X^−/−^ cells were X-gal stained at P5 following transduction with HDAd-*UbC-lacZ*-p*GGTA1* vector (GGTA1) in the presence or absence of enhancer treatment. Based on previous observations, non-integrated transgenes are largely lost by P5; therefore, we considered five passages sufficient to detect the differences in lacZ-positive cell percentages. Treatments were administered one day before transduction, refreshed during and immediately after transduction, and removed two days later (Fig. 6A). A non-targeting *lacZ* vector (NT) was included as a control for residual episomal *lacZ* expression at P5. Since all groups received the same vector dose, transduction efficiency was assumed to be comparable among treatments, and X-gal staining was not performed at P0. Preliminary screening identified M3814 as the most effective agent, with 4 μM producing the greatest enhancement of integration, whereas the remaining compounds had minimal effects (Fig. S2A). We therefore evaluated each compound in combination with M3814 (Fig. 6B). X-gal staining revealed that M3814 alone increased the lacZ-positive cell frequencies from 2.6% to 4.0% (Fig. 6C, D). Among all combinations tested, only NaBu + M3814 produced a statistically significant improvement, increasing integration efficiency to 6.4%, representing a 2.5-fold increase over the untreated control (Fig. 6C, D). These findings suggest that the combined NHEJ inhibition and chromatin remodeling can substantially enhance CRISPR/Cas9-mediated integration efficiency in PK15 W1282X^−/−^ cells. Given the 2.5-fold increase achieved with NaBu + M3814 treatment, we reasoned that combining this treatment with the dual-locus-targeting strategy may enable detectable *K18-CFTR* expression in PK15 W1282X^−/−^ cells.

**Figure 6.**
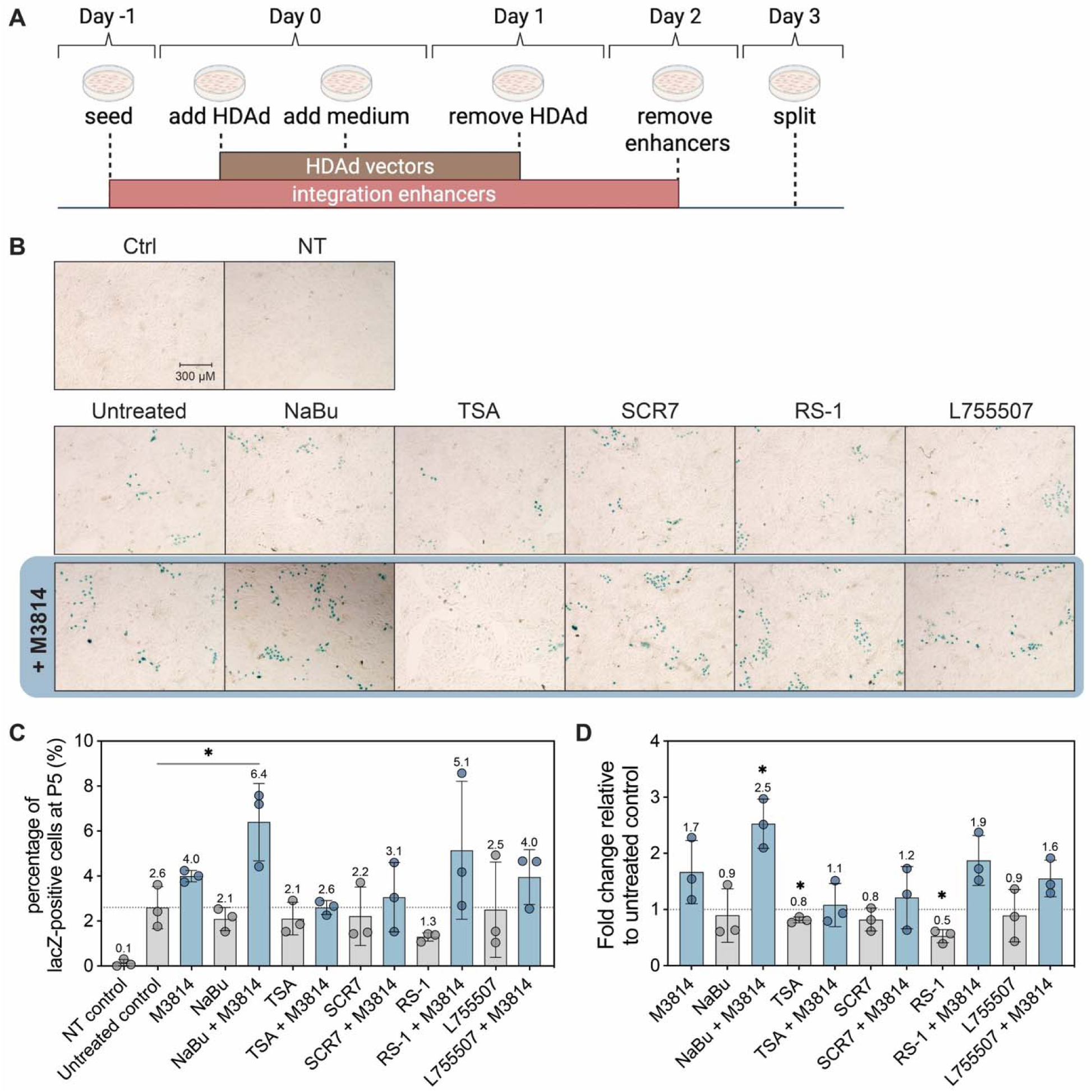
Integration-enhancing molecule screening in PK15 W1282X^−/−^ cell. **(A)** Schematic of HDAd delivery and enhancer treatment timeline. Cells were seeded in the presence of integration enhancers in growth medium. The next day, HDAd vectors were added to cells for 4 hours and were stopped with growth medium containing10% FBS. HDAd vectors were removed at 1 day post infection (dpi) by a medium change, and the enhancers were removed at 2 dpi. Cells were split to a new plate at 3 dpi. *Created with BioRender.com.* **(B)** X-gal staining images of PK15 W1282X^−/−^ cells at P5 after 30 MOI transduction of HDAd-*UbC-lacZ*-p*GGTA1* vector. Cells were also transduced with the NT vector as a negative control (image not shown). Integration enhancers were added one day before transduction and removed on day 2 post-transduction. The dose tested was 4 μM M3814, 400 μM Auxin, 2 mM NaBu, 300 nM TSA, 2 μM SCR7, 5 μM RS-1, and 5 μM L755507. **(C)** Percentages of lacZ-positive cells at P5. **(D)** Fold change of lacZ-positive percentage relative to untreated control. *Ordinary one-way ANOVA (B) and one-sample t test (C), *p < 0.05; n = 3*.

### CFTR replacement in PK15 W1282X^−/−^ cells can be improved with DEAE-dextran together with NaBu + M3814 treatment

To mitigate the negative effects of combined NaBu (2 mM) + M3814 (4 μM) treatment with high doses of HDAd vectors on cell growth and viability, the M3814 concentration was reduced to 3 μM. Additionally, a transduction enhancer was used to further reduce the required HDAd dose. DEAE-dextran is a positively charged polymer that facilitates viral uptake and enhances gene transfer efficiency, including for adenoviral and lentiviral vectors [43–45]. We previously applied DEAE-dextran to enhance HDAd transduction in human airway epithelial cells at 520.5 ng per 100 MOI and *in vivo* at 40 μg/mL [46, 47]. In the present study, a dose of 1041 ng per 100 MOI were used, enabling a reduction of HDAd dose to 50 MOI while still achieving WT-level CFTR channel function in PK15 W1282X^−/−^ cells (Fig. S2B, Fig. 7A). In comparison, 100 MOI was required in the absence of DEAE-dextran (Fig. 5H).

**Figure 7.**
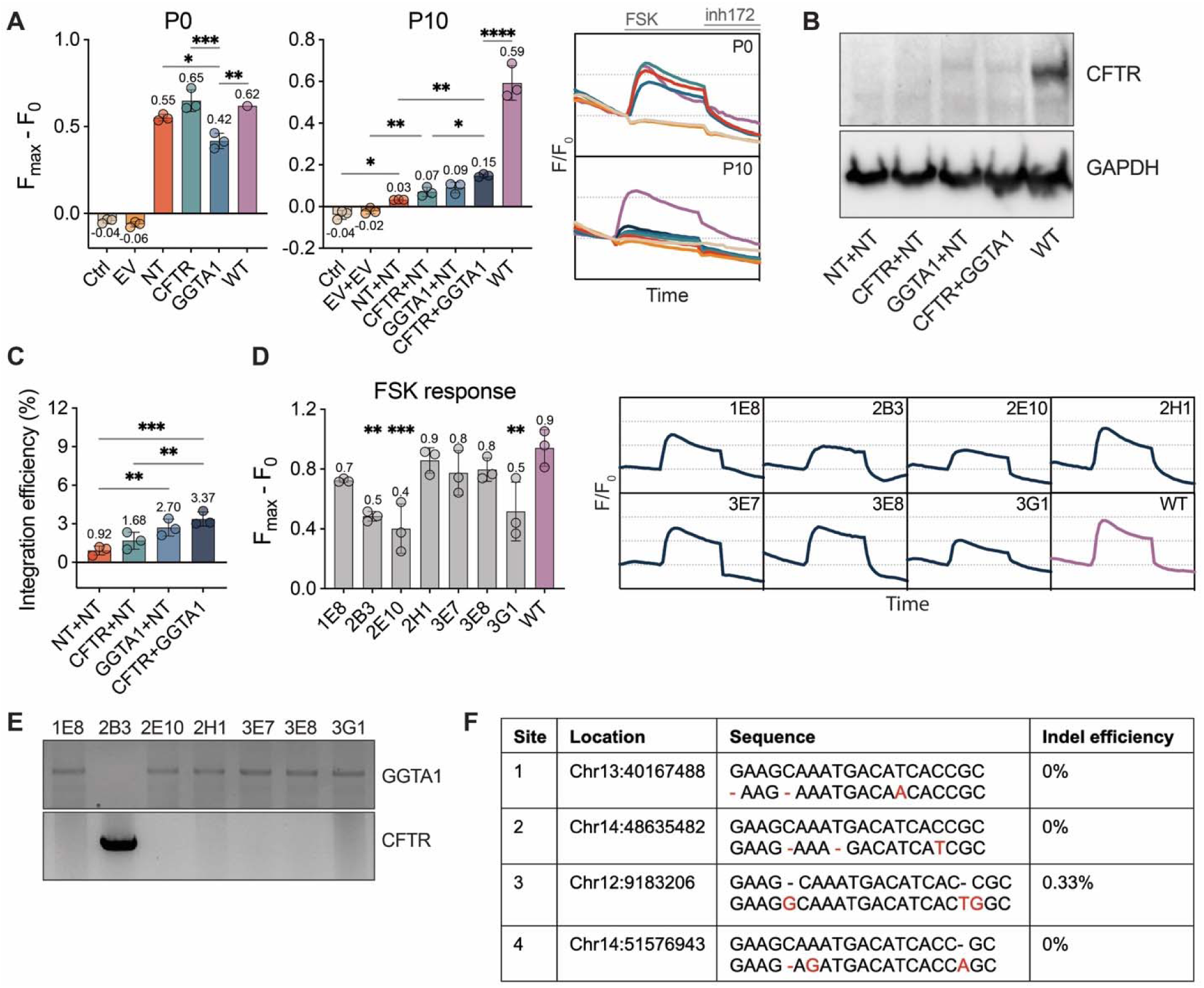
Restoration of CFTR expression in PK15 W1282X^−/−^ cells with integration enhancer treatment. PK15 W1282X^−/−^ cells were transduced with a total of 50 MOI HDAd vectors together with DEAE-dextran and NaBu+M3814 treatment. **(A)** CFTR function measured by FLIPR assay at P0 and P10. *Top panel:* change in baseline-normalized membrane fluorescence (F/F_0_) recorded over 27 min with forskolin (FSK) added at 5 min, and CFTRinh-172 (inh172) added at 15 min. *Bottom panel:* highest increase in fluorescence (F_max_-F_0_) after FSK activation. **(B)** Western blot of CFTR protein at P10 with GAPDH as the loading control. **(C)** Integration efficiency measured with ddPCR. **(D)** FLIPR assay showing 7 selected single-cell clones with channel function from the CFTR+GGTA1 group. CFTR+GGTA1 cells were single cell sorted to assess CFTR function. *Left panel:* highest increase in fluorescence (F_max_-F_0_) after FSK activation. *Right panel:* change in baseline-normalized membrane fluorescence (F/F_0_) recorded over 27 min with IBMX and forskolin (I&F) added at 5 min and CFTRinh-172 (inh172) added at 15 min. Statistical significance was determined relative to the WT control. (**E)** Junction PCR of the 7 clones to confirm the location of *K18-CFTR* integration at the *CFTR* or *GGTA1* locus. **(F)** Off-target analysis of four predicted sites for the *CFTR* target site. These regions were PCR amplified from the DNA of CFTR+GGTA1 cells, sequenced, and analyzed using the ICE tool for indel frequency. Percentages presented were averages of the triplicates. Ctrl, untransduced control; EV, empty vector; NT, non-targeting vector with *K18-CFTR* but not *Cas9*; *Two-way ANOVA*; *\*p < 0.05, **p < 0.01, ***p < 0.001, ****p < 0.0001; n = 3*.

Accordingly, PK15 W1282X^−/−^ cells were transduced with 50 MOI of HDAd vector at both P0 and P3 as described previously (Fig. 3A). DEAE-dextran (1041 ng per 100 MOI) was added during each transduction. NaBu (2 mM) + M3814 (3 μM) treatment were applied starting one day before transduction and removed two days after, with fresh compounds added during medium changes (Fig. 6A). FLIPR was performed at P10 when most episomal copies of *K18-CFTR* were expected to be lost (Fig. 7A). Under these conditions, CFTR+GGTA1 vectors produced the highest FSK-stimulated response, corresponding to 25.4% of WT activity (Fig. 7A). This response was significantly higher than CFTR+NT but not GGTA1+NT, consistent with the Western blot results at P10, where the CFTR protein was detected in GGTA1+NT and CFTR+GGTA1 (Fig. 7B). ddPCR analysis showed overall higher integration efficiencies compared with the previous experiment without enhancer treatments (Fig. 5E). *K18-CFTR* Integration in CFTR+GGTA1 cells was slightly higher than in single-locus-targeting groups, consistent with the trend observed in functional assays (Fig. 7A, C, CFTR+GGTA1 > GGTA1+NT > CFTR+NT). To assess CFTR function at the single cell level, CFTR+GGTA1 cells were sorted, and seven single cell clones with detectable CFTR activity were isolated (Fig. 7D). The CFTR function ranged from 42% to 91% of WT levels. Junction PCR revealed that *K18-CFTR* integration occurred predominantly at the *GGTA1* locus, with only one clone showing integration at the *CFTR* locus, suggesting higher efficiency of *GGTA1*-targeted integration in PK15 W1282X^−/−^ cells (Fig. 7E). These results are consistent with the observation that a 3.37% integration frequency can yield a substantially larger functional effect (25.4% of WT activity), indicating strong per allele functional impact of corrected cells (Fig. 7A, C).

Finally, off-target analysis of the CFTR-targeting vector was performed, as the GGTA1 vector has been previously examined and reported. Among the 46 predicted potential off-target sites with a distance of 3 from the target sequence, four were selected for sequencing, all located in non-coding regions. Amplicon sequencing of the CFTR+GGTA1 cells detected off-target editing at one site (0.33%), with no detectable editing at the remaining three sites (Fig. 7F, Fig. S3A, B) [28]. These values were lower than those previously observed for the GGTA1-targeting vector [22, 29].

Together, these results demonstrated that (i) NaBu + M3814 treatment during transduction improved the integration of *UbC-lacZ*, as well as *K18-CFTR*, in PK15 W1282X^−/−^ cells; (ii) the increased integration also benefited from DEAE-dextran treatment, which reduced the HDAd dose required for sufficient editing; (iii) the dual-locus-targeting method improves integration frequencies, with *GGTA1* serving as a more efficient integration site than *CFTR* in PK15 W1282X^−/−^ cells. (Fig. 7A, C).

## DISCUSSION

One of the main challenges with CRISPR/Cas9-mediated editing is low integration efficiency. Since editing outcomes depend on the repair of DSBs created by Cas9, the frequency of the desired edit is limited by the relatively rare HDR pathway, which is active only during the S or G2 phases of the cell cycle, whereas the constitutively active and error-prone NHEJ pathway leads predominantly to on-target indel mutations. As a result, most repaired DNA carries undesired indels, which also prevent retargeting with the same CRISPR/Cas9 system because target or PAM sequences are often disrupted following NHEJ repair. New Cas9-derived systems, including base-editing and prime-editing, have been developed to bypass the requirement for DSBs and HDR by using a Cas9 nickase [12–14]. Theoretically, these approaches may improve editing efficiency by avoiding DSB-induced NHEJ and enabling editing throughout the cell cycle. However, editing efficiency remains highly dependent on cell type, target locus, and DNA repair context. Therefore, for CF gene therapy, each mutation and model system still requires extensive optimization of the systems to achieve editing rates of clinical relevance. For example, a recent study used an early prime editing system (PE2) delivered by an HDAd and achieved only 2.4% correction of the W1282X *CFTR* mutation in human induced pluripotent stem cells [48]. Thus, the HDR-dependent CRISPR/Cas9 system remains advantageous in versatility and broad applicability, and this study aims to improve its editing efficiency in porcine epithelial cell models for potential *in vivo* application.

Due to indel mutations generated by the dominant NHEJ pathway, most target or PAM sequences are altered and become untargetable. This not only produces undesired edits but also limits repeated targeting. To circumvent this limitation, we delivered two gene-editing cassettes targeting the porcine *CFTR* or *GGTA1* locus to increase the number of targetable sites and enhance overall integration. In our initial simultaneous HDAd delivery approach, integration was detected at both loci, but with limited improvement in total efficiency compared to single-locus targeting. We reasoned that the fraction of cells in S/G2 phase during transduction is limiting and cannot be further increased by higher vector dose. Therefore, to increase the number of HDR-competent cells, we applied sequential transduction, assuming that cells in HDR-permissive phases during the second delivery are partially distinct from those targeted during the first. Consistent with this hypothesis, sequential delivery significantly increased *UbC-lacZ* integration in IPEC-J2 cells, reaching 16.5% when CFTR and GGTA1 vectors were delivered separately, with a three-passage interval (Fig. 2E). In PK15 cells, which exhibited lower overall integration efficiency, CFTR+GGTA1 and GGTA1+CFTR groups showed similar integration levels; however, only the former reached statistical significance due to variability (Fig. 2G). The low efficiencies in PK15 cells are consistent with their lower proliferation rate compared with intestinal epithelial cells, resulting in fewer cells in HDR-permissive phases [49, 50]. For the *K18-CFTR* transgene, CFTR+GGTA1 in WT IPEC-J2 cells achieved 6.5% integration (Fig. 3G), without a detectable increase in CFTR expression over endogenous levels (Fig. 3H-J). Since the *CFTR* target site is located within intron 1, NHEJ-induced indels may not affect the native CFTR expression, but HDR-mediated integration of *K18-CFTR* disrupts the endogenous WT allele. Thus, the 4.7% integration observed at the *CFTR* locus in CFTR+NT also represents a proportional loss of endogenous *CFTR* alleles. Consequently, an increase in *CFTR* copy number relies primarily on *GGTA1* locus integration, explaining the lack of detectable functional change in CFTR+GGTA1 groups (Fig. 3G-J). To better capture transgene-derived CFTR expression, a PK15 W1282X^−/−^ model was generated. Due to low transduction and integration efficiencies in PK15 cells, enhancers were applied together with the dual-locus-targeting method. Under these conditions, CFTR+GGTA1 achieved 3.4%, resulting in 25.4% of WT CFTR activity and detectable CFTR protein expression, which was not observed previously with the GGTA1 vector in knock-out cells (Fig. 7A-C) [22]. Single-cell analysis further confirmed that clones with single-locus *K18-CFTR* integration exhibited sufficient CFTR activity (Fig. 7D, E). Overall, we demonstrated that the dual-locus-targeting strategy, combined with the sequential delivery, enhanced transgene integration in IPEC-J2 cells and in PK15 cells with enhancer treatment.

We found that integration efficiency can be enhanced by combining the NHEJ inhibitor M3814 with the HDACi NaBu (Fig. 6C, D). M3814 inhibits DNA-dependent protein kinase (DNA-PK), a key component of the Ku70/Ku80-DNA-PKcs complex that keeps the DNA ends in proximity for efficient NHEJ [36, 51]. As a result, M3814 (Peposertib) has also been studied as an anti-cancer agent by inhibiting DNA repair in tumour cells [52, 53]. Prolonged exposure to M3814 may therefore compromise genome stability. HDACi have also been reported to influence DNA repair pathways. NaBu has been shown to enhance Cas9 cutting efficiency, whereas TSA can improve both NHEJ and HDR frequencies [54, 55]. Although we did not directly assess these mechanisms, it is plausible that combined Cas9 enhancement and NHEJ suppression contributed to increased knock-in efficiency in PK15 cells. Further investigation is needed to evaluate translational applicability *in vivo*.

While this study provided insights into the circumvention of DSB-associated low transgene integration for CRISPR/Cas9, several limitations should be acknowledged. First, optimal editing typically requires screening multiple gRNAs, but only a single gRNA per locus was tested due to the labour-intensive development and production of HDAd vectors. gRNA performance may vary in different cell types. Therefore, effective gRNA design also needs cell type-specific optimization to enable efficient editing. Secondly, an ideal CF airway epithelial model was not available, necessitating the use of PK15 W1282X^−/−^ cells, which exhibited lower transduction efficiency. The lower integration efficiencies in PK15 cells compared to IPEC-J2 cells were consistent with those observed with the *UbC-lacZ* donor gene (Fig. 2G). The highest integration efficiency observed was 16.5% and 6.5% in IPEC-J2 cells, but 5.9% and below 2% in PK15 cells for *UbC-lacZ* and *K18-CFTR*, respectively (Fig. 2E, G; Fig. 3G; Fig. 5E). Since integration efficiencies of *UbC-lacZ* and *K18-CFTR* were measured using different methods, the values are not directly comparable. Intestinal epithelial cells undergo faster turnover, whereas kidney epithelial cells tend to remain quiescent unless injured, which may explain the lower and more difficult-to-enhance integration efficiencies in PK15 compared to IPEC-J2 cells [49, 50]. With DEAE-dextran and NaBu+M3814 treatment to enhance transduction and integration, CFTR+GGTA1 achieved 3.37% *K18-CFTR* integration in PK15 mutant cells, significantly higher than cells receiving CFTR+NT (Fig. 7C). In comparison to recent studies that used CRISPR/Cas9 in human airway epithelial cells, the integration efficiencies were 4-11% for CFTR cDNA without enrichment using AAV vectors [56, 57]. CF porcine airway epithelial cells with faster turnover due to persistent inflammation may produce more efficient *CFTR* KI than kidney cells. Additionally, applying the dual-locus-targeting method in human airway epithelial cells using their optimized CRISPR/Cas9 systems may further enhance the integration efficiency above the reported 4-11%.

Additional factors influencing HDR efficiency include the size of the donor gene and homology arms and accessibility to the repair template. The large *K18-CFTR* cassette (8.4 kb) likely reduced integration efficiency compared with *UbC-lacZ* (6 kb), consistent with previous reports that KI efficiency is significantly decreased when the insert is increased from 1.23 kb to 1.80 kb [58]. Since larger donor genes also necessitate longer homology arms (HA) for efficient recombination, the HAs of our HDAd vectors for both targeting loci were designed to be around 3 kb in length. However, long HAs also increase cloning difficulty [59–61], as well as the risk of off-target integration due to partial homology elsewhere [62]. One study demonstrated improved integration efficiency (7.54%) of a 20 kb transgene in porcine fetal fibroblasts with one short arm (0.3 kb) and one long arm (3.8 kb) [63]. Therefore, editing outcomes may be further improved with HAs optimization. Next, availability to repair templates is also pivotal to efficient HDR. HDAd vectors were designed to carry all components in a single vector to ensure that transduced cells receive everything required for gene editing rather than partial elements, but it also prevents the adjustment of the Cas9-to-repair template ratio. Accumulating evidence supports that improved editing efficiency is associated with a higher ratio of donor template or proximity of the template by tethering to Cas9 [64–67]. Overall, these factors can be further investigated in future optimization studies.

Finally, adding a target locus may increase the frequencies of off-target edits and on-target indel mutations. A recent study investigated the safety of full-length *CFTR* cDNA insertion at human *CFTR* exon 1 and detected 0.01% allele translocation, mostly in non-oncogenic sites, and no chromosomal rearrangements [68]. We used the ICE tool to analyze the Sanger sequencing result of four predicted off-target sites for the *CFTR* locus and observed indels at only one site with a frequency of 0.33% (Fig. 7F). The off-target effects of the GGTA1 vector have been reported previously to be mostly below 1% using the same method and were therefore not repeated here [22]. Note that the 0% edit frequency reported here indicates frequencies too low to be detected by Sanger sequencing (Fig. 7F). However, it can be concluded that the dual-locus-targeting method does produce more undesired edits, with an increased number of potential off-target editing sites, but most are detected at low frequencies. While off-target effects could be reduced by using high-fidelity Cas9 variants [69–71], the undesired on-target indels can potentially be addressed using the “Double Tap” [72] and the “Recursive Editing” [73] approaches that involve extra gRNAs for retargeting. However, this method also carries an increasing risk of off-target edits for each extra gRNA. Enriching corrected cell populations has also been frequently used to avoid undesired editing outcomes, which is commonly used for *ex vivo* cell therapy, as *in vivo* selection is still challenging to perform safely, especially for CF, where corrected cells and mutant cells are equally viable [74–76].

Immune responses to HDAd vectors also remain a major challenge. Ongoing studies are exploring potential approaches, which will be further examined in future studies. We have previously shown that cyclophosphamide reduces the expression of T cell genes and the level of anti-adenoviral antibodies in the mouse airway following repeated HDAd delivery [47]. Immune reaction against adenoviral vectors can also be suppressed by exogenously expressing the anti-inflammatory cytokine IL-10 [77–79]. These strategies may be combined with other common immune evasion approaches, such as vector shielding with polyethylene glycol (PEG) and fibre and capsid engineering [80–83].

Taken together, we demonstrated a simple method of targeting different loci sequentially to overcome the integration limitation of CRISPR/Cas9. The results of this study enhance our understanding of the HDR-mediated integration mechanism, which may guide the optimization of future gene editing experiments. Using the dual-locus-targeting strategy, the highest integration efficiency (16.5%) of the *UbC-lacZ* transgene was achieved with the HDAd-delivered CRISPR/Cas9 system. This efficiency may lead to detectable CFTR expression in mutant cells transduced with HDAd vectors with the *CFTR* transgene. These findings highlight potential strategies to improve the efficacy of CRISPR/Cas9-mediated transgene integration. Incorporating these strategies with upgraded Cas9 systems and optimized gRNA design is expected to yield further improved outcomes, supporting the feasibility of achieving a clinically relevant level of CFTR rescue in CF cells.

## ACKNOWLEDGEMENT

We acknowledge Dr. Ranmal Avinash Bandara for valuable discussions and suggestions regarding the strategy for generating the homozygous mutant cell line.

## AUTHOR CONTRIBUTIONS

Ziyan Rachel Chen: Formal analysis, Investigation, Methodology, Writing-original draft, Visualization. Zhichang Peter Zhou: Resources, Validation, Writing-review & editing. Rongqi Cathleen Duan: Resources, Methodology, Writing-review & editing. Amy Wong: Funding Acquisition, Writing-review & editing. Hartmut Grasemann: Writing-review & editing. Christine Bear: Writing-review & editing. Jim Hu: Conceptualization, Methodology, Supervision, Funding Acquisition, Writing-review & editing.

## SUPPLEMENTARY DATA STATEMENT

Supplementary Data are available online.

## DATA AVAILABILITY

The data underlying this article are available in the article and in its online supplementary material.

## REFERENCES

[1] Kerem B, Rommens JM, Buchanan JA, et al. Identification of the cystic fibrosis gene: genetic analysis. Science 1989; 245: 1073–1080.

[2] Riordan JR, Rommens JM, Kerem B, et al. Identification of the cystic fibrosis gene: cloning and characterization of complementary DNA. Science 1989; 245: 1066–1073.

[3] Rommens JM, Iannuzzi MC, Kerem B-S, et al. Identification of the Cystic Fibrosis Gene: Chromosome Walking and Jumping. Science 1989; 245: 1059–1065.

[4] Brodlie M, Haq IJ, Roberts K, et al. Targeted therapies to improve CFTR function in cystic fibrosis. Genome Med 2015; 7: 101.

[5] Pranke I, Golec A, Hinzpeter A, et al. Emerging Therapeutic Approaches for Cystic Fibrosis. From Gene Editing to Personalized Medicine. Front Pharmacol 2019; 10: 121.

[6] Grasemann H, Ratjen F. Cystic Fibrosis. N Engl J Med 2023; 389: 1693–1707.

[7] Lopez A, Daly C, Vega-Hernandez G, et al. Elexacaftor/tezacaftor/ivacaftor projected survival and long-term health outcomes in people with cystic fibrosis homozygous for *F508del*. Journal of Cystic Fibrosis 2023; 22: 607–614.

[8] Koehler DR, Sajjan U, Chow Y-H, et al. Protection of Cftr knockout mice from acute lung infection by a helper-dependent adenoviral vector expressing Cftr in airway epithelia. Proc Natl Acad Sci U S A 2003; 100: 15364–15369.

[9] Toietta G, Koehler DR, Finegold MJ, et al. Reduced inflammation and improved airway expression using Helper-Dependent adenoviral vectors with a k18 promoter. Molecular Therapy 2003; 7: 649–658.

[10] Sl F, Ph K, P N, et al. Gene transfer of CFTR to airway epithelia: low levels of expression are sufficient to correct Cl-transport and overexpression can generate basolateral CFTR. American journal of physiology Lung cellular and molecular physiology; 289. Epub ahead of print December 2005. DOI: 10.1152/ajplung.00049.2005.

[11] Ran FA, Hsu PD, Wright J, et al. Genome engineering using the CRISPR-Cas9 system. Nat Protoc 2013; 8: 2281–2308.

[12] Komor AC, Kim YB, Packer MS, et al. Programmable editing of a target base in genomic DNA without double-stranded DNA cleavage. Nature 2016; 533: 420–424.

[13] Gaudelli NM, Komor AC, Rees HA, et al. Programmable base editing of A•T to G•C in genomic DNA without DNA cleavage. Nature 2017; 551: 464–471.

[14] Anzalone AV, Randolph PB, Davis JR, et al. Search-and-replace genome editing without double-strand breaks or donor DNA. Nature 2019; 576: 149–157.

[15] Jinek M, Chylinski K, Fonfara I, et al. A programmable dual-RNA-guided DNA endonuclease in adaptive bacterial immunity. Science 2012; 337: 816–821.

[16] Bolderson E, Tomimatsu N, Richard DJ, et al. Phosphorylation of Exo1 modulates homologous recombination repair of DNA double-strand breaks. Nucleic Acids Res 2010; 38: 1821–1831.

[17] Welsh MJ, Rogers CS, Stoltz DA, et al. Development of a Porcine Model of Cystic Fibrosis. Trans Am Clin Climatol Assoc 2009; 120: 149–162.

[18] Chow YH, O’Brodovich H, Plumb J, et al. Development of an epithelium-specific expression cassette with human DNA regulatory elements for transgene expression in lung airways. Proc Natl Acad Sci U S A 1997; 94: 14695–14700.

[19] Chow YH, Plumb J, Wen Y, et al. Targeting transgene expression to airway epithelia and submucosal glands, prominent sites of human CFTR expression. Mol Ther 2000; 2: 359–367.

[20] Cao H, Machuca TN, Yeung JC, et al. Efficient gene delivery to pig airway epithelia and submucosal glands using helper-dependent adenoviral vectors. Mol Ther Nucleic Acids 2013; 2: e127.

[21] Cao H, Ouyang H, Ip W, et al. Testing gene therapy vectors in human primary nasal epithelial cultures. Mol Ther Methods Clin Dev 2015; 2: 15034.

[22] Zhou ZP, Yang LL, Cao H, et al. In Vitro Validation of a CRISPR-Mediated CFTR Correction Strategy for Preclinical Translation in Pigs. Hum Gene Ther 2019; 30: 1101–1116.

[23] Bartoszewski R, Rab A, Jurkuvenaite A, et al. Activation of the Unfolded Protein Response by ΔF508 CFTR. Am J Respir Cell Mol Biol 2008; 39: 448–457.

[24] Bartoszewski R, Rab A, Fu L, et al. CFTR expression regulation by the unfolded protein response. Methods Enzymol 2011; 491: 3–24.

[25] Sandig V, Youil R, Bett AJ, et al. Optimization of the helper-dependent adenovirus system for production and potency in vivo. Proceedings of the National Academy of Sciences 2000; 97: 1002–1007.

[26] Palmer DJ, Ng P. Methods for the production of helper-dependent adenoviral vectors. Methods Mol Biol 2008; 433: 33–53.

[27] Koehler DR, Sajjan U, Chow Y-H, et al. Protection of Cftr knockout mice from acute lung infection by a helper-dependent adenoviral vector expressing Cftr in airway epithelia. Proc Natl Acad Sci U S A 2003; 100: 15364–15369.

[28] Labun K, Montague TG, Krause M, et al. CHOPCHOP v3: expanding the CRISPR web toolbox beyond genome editing. Nucleic Acids Res 2019; 47: W171–W174.

[29] Hsiau T, Conant D, Rossi N, et al. Inference of CRISPR Edits from Sanger Trace Data. 2019; 251082.

[30] Chang HHY, Watanabe G, Gerodimos CA, et al. Different DNA End Configurations Dictate Which NHEJ Components Are Most Important for Joining Efficiency. J Biol Chem 2016; 291: 24377–24389.

[31] Yu C, Liu Y, Ma T, et al. Small molecules enhance CRISPR genome editing in pluripotent stem cells. Cell Stem Cell 2015; 16: 142–147.

[32] Li G, Zhang X, Zhong C, et al. Small molecules enhance CRISPR/Cas9-mediated homology-directed genome editing in primary cells. Sci Rep 2017; 7: 8943.

[33] Jayathilaka K, Sheridan SD, Bold TD, et al. A chemical compound that stimulates the human homologous recombination protein RAD51. Proc Natl Acad Sci U S A 2008; 105: 15848–15853.

[34] Song J, Yang D, Xu J, et al. RS-1 enhances CRISPR/Cas9- and TALEN-mediated knock-in efficiency. Nat Commun 2016; 7: 10548.

[35] Harnor SJ, Brennan A, Cano C. Targeting DNA-Dependent Protein Kinase for Cancer Therapy. ChemMedChem 2017; 12: 895–900.

[36] Riesenberg S, Chintalapati M, Macak D, et al. Simultaneous precise editing of multiple genes in human cells. Nucleic Acids Research 2019; 47: e116.

[37] Riesenberg S, Kanis P, Macak D, et al. Efficient high-precision homology-directed repair-dependent genome editing by HDRobust. Nat Methods 2023; 1–12.

[38] Srivastava M, Nambiar M, Sharma S, et al. An inhibitor of nonhomologous end-joining abrogates double-strand break repair and impedes cancer progression. Cell 2012; 151: 1474–1487.

[39] Maruyama T, Dougan SK, Truttmann MC, et al. Increasing the efficiency of precise genome editing with CRISPR-Cas9 by inhibition of nonhomologous end joining. Nat Biotechnol 2015; 33: 538–542.

[40] Davie JR. Inhibition of histone deacetylase activity by butyrate. J Nutr 2003; 133: 2485S–2493S.

[41] Yoshida M, Horinouchi S, Beppu T. Trichostatin A and trapoxin: novel chemical probes for the role of histone acetylation in chromatin structure and function. Bioessays 1995; 17: 423–430.

[42] Vigushin DM, Ali S, Pace PE, et al. Trichostatin A is a histone deacetylase inhibitor with potent antitumor activity against breast cancer in vivo. Clin Cancer Res 2001; 7: 971–976.

[43] Graham FL, van der Eb AJ. A new technique for the assay of infectivity of human adenovirus 5 DNA. Virology 1973; 52: 456–467.

[44] Gregory LG, Harbottle RP, Lawrence L, et al. Enhancement of adenovirus-mediated gene transfer to the airways by DEAE dextran and sodium caprate in vivo. Mol Ther 2003; 7: 19–26.

[45] Amadeo F, Hanson V, Murray P, et al. DEAE-Dextran Enhances the Lentiviral Transduction of Primary Human Mesenchymal Stromal Cells from All Major Tissue Sources Without Affecting Their Proliferation and Phenotype. Mol Biotechnol 2023; 65: 544–555.

[46] Lee C-M, Gupta S, Wang J, et al. Epithelium-specific Ets transcription factor-1 acts as a negative regulator of cyclooxygenase-2 in human rheumatoid arthritis synovial fibroblasts. Cell Biosci 2016; 6: 43.

[47] Cao H, Duan R, Hu J. Overcoming Immunological Challenges to Helper-Dependent Adenoviral Vector-Mediated Long-Term CFTR Expression in Mouse Airways. Genes (Basel) 2020; 11: 565.

[48] Li C, Liu Z, Anderson J, et al. Prime editing-mediated correction of the CFTR W1282X mutation in iPSCs and derived airway epithelial cells. PLoS One 2023; 18: e0295009.

[49] Nadasdy T, Laszik Z, Blick KE, et al. Proliferative activity of intrinsic cell populations in the normal human kidney. J Am Soc Nephrol 1994; 4: 2032–2039.

[50] Darwich AS, Aslam U, Ashcroft DM, et al. Meta-Analysis of the Turnover of Intestinal Epithelia in Preclinical Animal Species and Humans. Drug Metabolism and Disposition 2014; 42: 2016–2022.

[51] Liang S, Blundell TL. Human DNA-dependent protein kinase activation mechanism. Nat Struct Mol Biol 2023; 30: 140–147.

[52] Zenke FT, Zimmermann A, Sirrenberg C, et al. Pharmacologic Inhibitor of DNA-PK, M3814, Potentiates Radiotherapy and Regresses Human Tumors in Mouse Models. Mol Cancer Ther 2020; 19: 1091–1101.

[53] van Bussel MTJ, Awada A, de Jonge MJA, et al. A first-in-man phase 1 study of the DNA-dependent protein kinase inhibitor peposertib (formerly M3814) in patients with advanced solid tumours. Br J Cancer 2021; 124: 728–735.

[54] Li G, Zhang X, Wang H, et al. Increasing CRISPR/Cas9-mediated homology-directed DNA repair by histone deacetylase inhibitors. The International Journal of Biochemistry & Cell Biology 2020; 125: 105790.

[55] Björnson Y, Huang CY, Rollins JL, et al. The effect of histone deacetylase inhibitors on the efficiency of the CRISPR/Cas9 system. Biochem Biophys Rep 2023; 35: 101513.

[56] Vaidyanathan S, Baik R, Chen L, et al. Targeted replacement of full-length CFTR in human airway stem cells by CRISPR-Cas9 for pan-mutation correction in the endogenous locus. Molecular Therapy 2022; 30: 223–237.

[57] Stack JT, Rayner RE, Nouri R, et al. DNA-PKcs inhibition improves sequential gene insertion of the full-length *CFTR* cDNA in airway stem cells. Molecular Therapy Nucleic Acids 2024; 35: 102339.

[58] Sakurai T, Takei N, Wei Y, et al. Efficient genome editing of two-cell mouse embryos via modified CRISPR/Cas electroporation. Sci Rep 2024; 14: 30347.

[59] Baker O, Tsurkan S, Fu J, et al. The contribution of homology arms to nuclease-assisted genome engineering. Nucleic Acids Res 2017; 45: 8105–8115.

[60] Paix A, Folkmann A, Goldman DH, et al. Precision genome editing using synthesis-dependent repair of Cas9-induced DNA breaks. Proceedings of the National Academy of Sciences 2017; 114: E10745–E10754.

[61] Zhang J-P, Li X-L, Li G-H, et al. Efficient precise knockin with a double cut HDR donor after CRISPR/Cas9-mediated double-stranded DNA cleavage. Genome Biol 2017; 18: 35.

[62] Ishii A, Kurosawa A, Saito S, et al. Analysis of the Role of Homology Arms in Gene-Targeting Vectors in Human Cells. PLoS One 2014; 9: e108236.

[63] Li G, Zhang X, Wang H, et al. CRISPR/Cas9-Mediated Integration of Large Transgene into Pig CEP112 Locus. G3 Genes|Genomes|Genetics 2020; 10: 467–473.

[64] Savic N, Ringnalda FC, Lindsay H, et al. Covalent linkage of the DNA repair template to the CRISPR-Cas9 nuclease enhances homology-directed repair. eLife 2018; 7: e33761.

[65] Aird EJ, Lovendahl KN, St. Martin A, et al. Increasing Cas9-mediated homology-directed repair efficiency through covalent tethering of DNA repair template. Commun Biol 2018; 1: 54.

[66] Shin SW, Kim D, Lee JS, et al. Controlling Ratios of Plasmid-Based Double Cut Donor and CRISPR/Cas9 Components to Enhance Targeted Integration of Transgenes in Chinese Hamster Ovary Cells. International Journal of Molecular Sciences; 22. Epub ahead of print 26 February 2021. DOI: 10.3390/ijms22052407.

[67] Zhou Z, Xiao J, Yin S, et al. Cas9-Rep fusion tethers donor DNA in vivo and boosts the efficiency of HDR-mediated genome editing. Plant Biotechnol J 2025; 23: 2006–2017.

[68] Vaidyanathan S, Kerschner JL, Paranjapye A, et al. Investigating adverse genomic and regulatory changes caused by replacement of the full-length CFTR cDNA using Cas9 and AAV. Molecular Therapy Nucleic Acids; 35. Epub ahead of print 12 March 2024. DOI: 10.1016/j.omtn.2024.102134.

[69] Kleinstiver BP, Pattanayak V, Prew MS, et al. High-fidelity CRISPR–Cas9 nucleases with no detectable genome-wide off-target effects. Nature 2016; 529: 490–495.

[70] Vakulskas CA, Dever DP, Rettig GR, et al. A high-fidelity Cas9 mutant delivered as a ribonucleoprotein complex enables efficient gene editing in human hematopoietic stem and progenitor cells. Nat Med 2018; 24: 1216–1224.

[71] Pedrazzoli E, Bianchi A, Umbach A, et al. An optimized SpCas9 high-fidelity variant for direct protein delivery. Molecular Therapy 2023; S1525001623001284.

[72] Bodai Z, Bishop AL, Gantz VM, et al. Targeting double-strand break indel byproducts with secondary guide RNAs improves Cas9 HDR-mediated genome editing efficiencies. Nat Commun 2022; 13: 2351.

[73] Möller L, Aird EJ, Schröder MS, et al. Recursive Editing improves homology-directed repair through retargeting of undesired outcomes. Nat Commun 2022; 13: 4550.

[74] Vaidyanathan S, Baik R, Chen L, et al. Targeted replacement of full-length CFTR in human airway stem cells by CRISPR-Cas9 for pan-mutation correction in the endogenous locus. Mol Ther 2022; 30: 223–237.

[75] Suzuki S, Sargent RG, Illek B, et al. TALENs Facilitate Single-step Seamless SDF Correction of F508del CFTR in Airway Epithelial Submucosal Gland Cell-derived CF-iPSCs. Molecular Therapy Nucleic Acids; 5. Epub ahead of print 1 January 2016. DOI: 10.1038/mtna.2015.43.

[76] Yan Z, Vorhies K, Feng Z, et al. Recombinant Adeno-Associated Virus-Mediated Editing of the G551D Cystic Fibrosis Transmembrane Conductance Regulator Mutation in Ferret Airway Basal Cells. Hum Gene Ther 2022; 33: 1023–1036.

[77] Kushwah R, Oliver JR, Duan R, et al. Induction of Immunological Tolerance to Adenoviral Vectors by Using a Novel Dendritic Cell-Based Strategy. Journal of Virology 2012; 86: 3422–3435.

[78] Yeung JC, Wagnetz D, Cypel M, et al. Ex Vivo Adenoviral Vector Gene Delivery Results in Decreased Vector-associated Inflammation Pre- and Post–lung Transplantation in the Pig. Molecular Therapy 2012; 20: 1204–1211.

[79] Guo Z, He Q, Zhang Y, et al. Mechanisms of Interleukin-10-Mediated Immunosuppression in Viral Infections. Pathogens 2025; 14: 989.

[80] Li H-J, Everts M, Pereboeva L, et al. Adenovirus tumor targeting and hepatic untargeting by a coxsackie/adenovirus receptor ectodomain anti-carcinoembryonic antigen bispecific adapter. Cancer Res 2007; 67: 5354–5361.

[81] Georgakopoulou A, Wang H, Kim J, et al. *In vivo* HSC transduction in humanized mice mediated by novel capsid-modified HDAd vectors. Molecular Therapy Methods & Clinical Development 2025; 33: 101448.

[82] Chillón M, Lee JH, Fasbender A, et al. Adenovirus complexed with polyethylene glycol and cationic lipid is shielded from neutralizing antibodies in vitro. Gene Ther 1998; 5: 995–1002.

[83] Schmid M, Ernst P, Honegger A, et al. Adenoviral vector with shield and adapter increases tumor specificity and escapes liver and immune control. Nat Commun 2018; 9: 450.

