## supplementary information for "A Dual-Locus-Targeting Strategy to Enhance CRISPR/Cas9-mediated CFTR Replacement via Helper-Dependent Adenoviral vector in porcine genome"

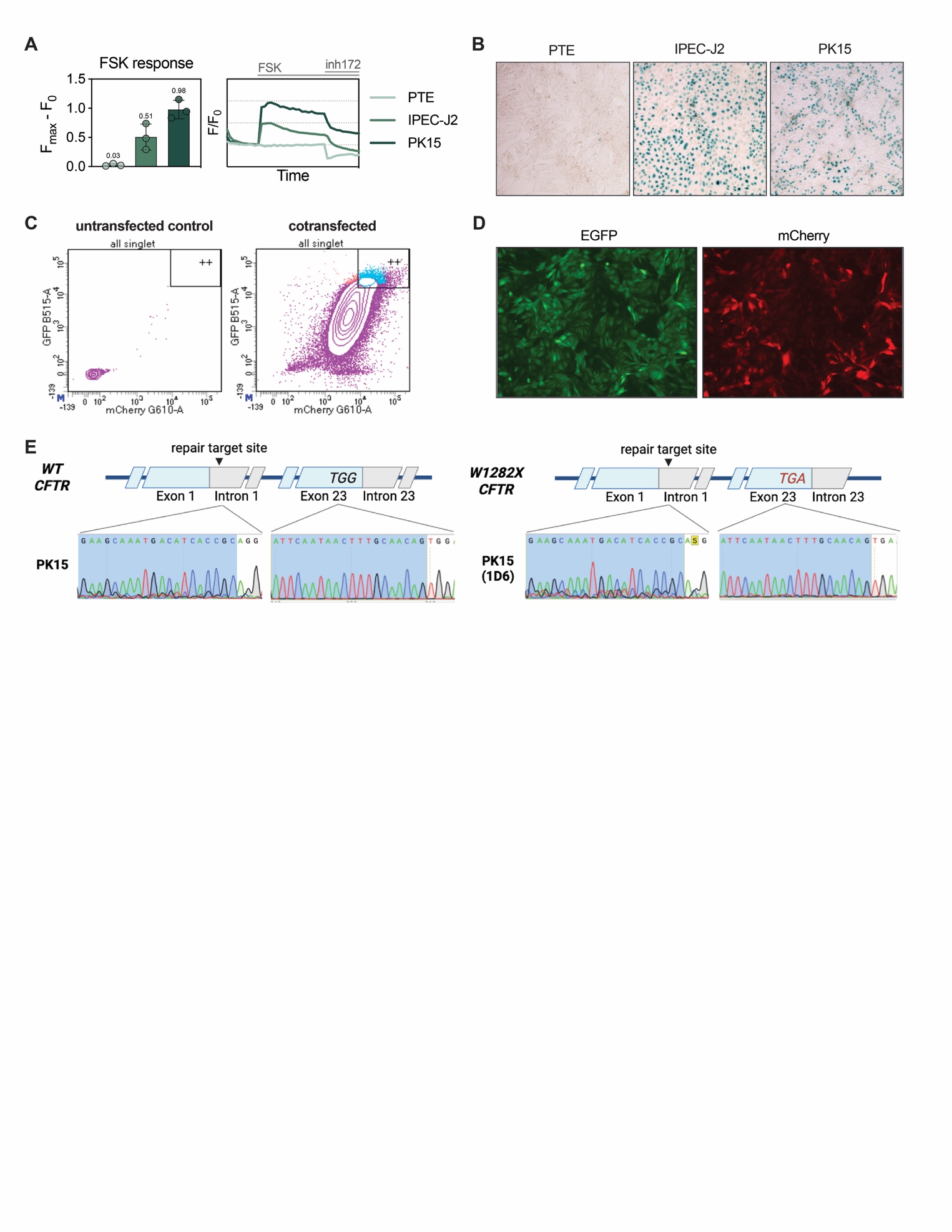

**Supplementary Figure 1. PK15 W1282X^-/-^ cell line generation. (A)** CFTR channel activity in the three candidate cell lines. PTE, porcine tracheal epithelial cell line. *Left panel:* highest increase in fluorescence (F_max_-F_0_) after forskolin (FSK) activation. *Right panel:* change in baseline-normalized membrane fluorescence (F/F_0_) recorded over 27 min with FSK added at 5 min and CFTRinh-172 (inh172) added at 15 min. *n = 1*. **(B)** X-gal staining of the three candidate cell lines following 25 MOI *UbC-lacZ* transduction to compare the transduction efficiency. Images were taken at two days post-delivery. *n = 1*. **(C)** mCherry^+^/EGFP^+^ gate for sorting in PK15 cells untransfected or cotransfected with the two plasmids designed for W1282X^-/-^ mutagenesis. **(D)** mCherry^+^/EGFP^+^ PK15 cells after the first selection step. **(E)** Sequencing of the W1282X mutation site at exon 23 and the subsequent *K18-CFTR* integration target site at intron 1 for both PK15 and the mutant clone PK15 (1D6) cells*.* *Created with BioRender.com.*

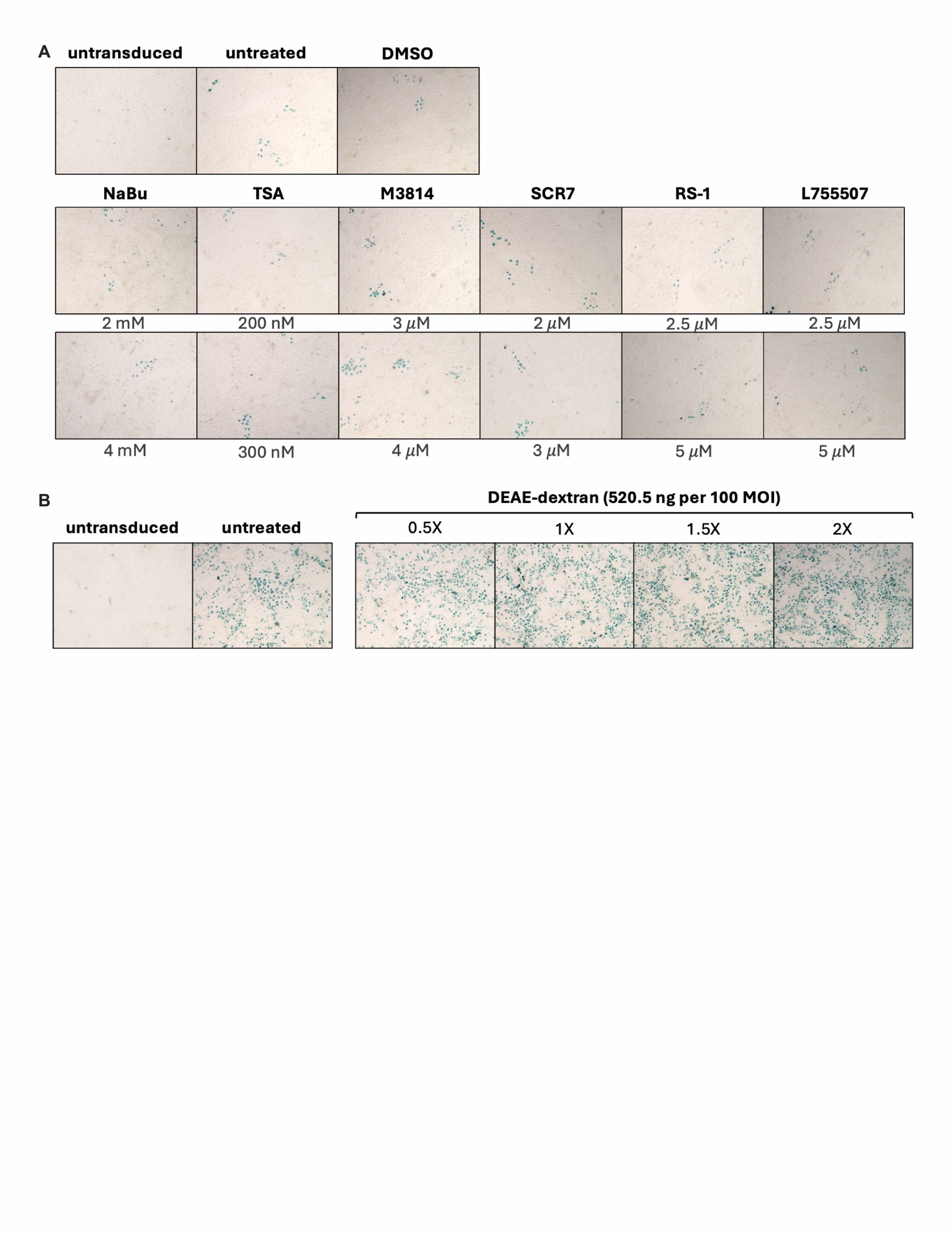

**Supplementary Figure 2. Preliminary tests for transduction and integration enhancers using *UbC-lacZ* vectors in PK15 W1282X^-/-^ cells. (A)** Cells were transduced with 10 MOI of non-targeting *UbC-lacZ* vector. HDAd viral particles were incubated with DEAE-dextran for 30 min at 37°C before adding to the cells. X-gal staining was performed at day 2 post-transduction to assess transduction efficiency. Increasing doses of DEAE-dextran was examined, with “1$\times$” representing the previously described dose of 520.5 ng per 100 MOI and 2$\times$ representing a two-fold increase. **(B)** Cells were treated with small molecules as described in Fig. 3.8A and transduced with 30 MOI of GGTA1-targeting *UbC-lacZ* vector. X-gal staining was performed at passage 5. DMSO was the vehicle control for TSA, M3814, SCR7, RS-1, and L755507. *n = 2.*

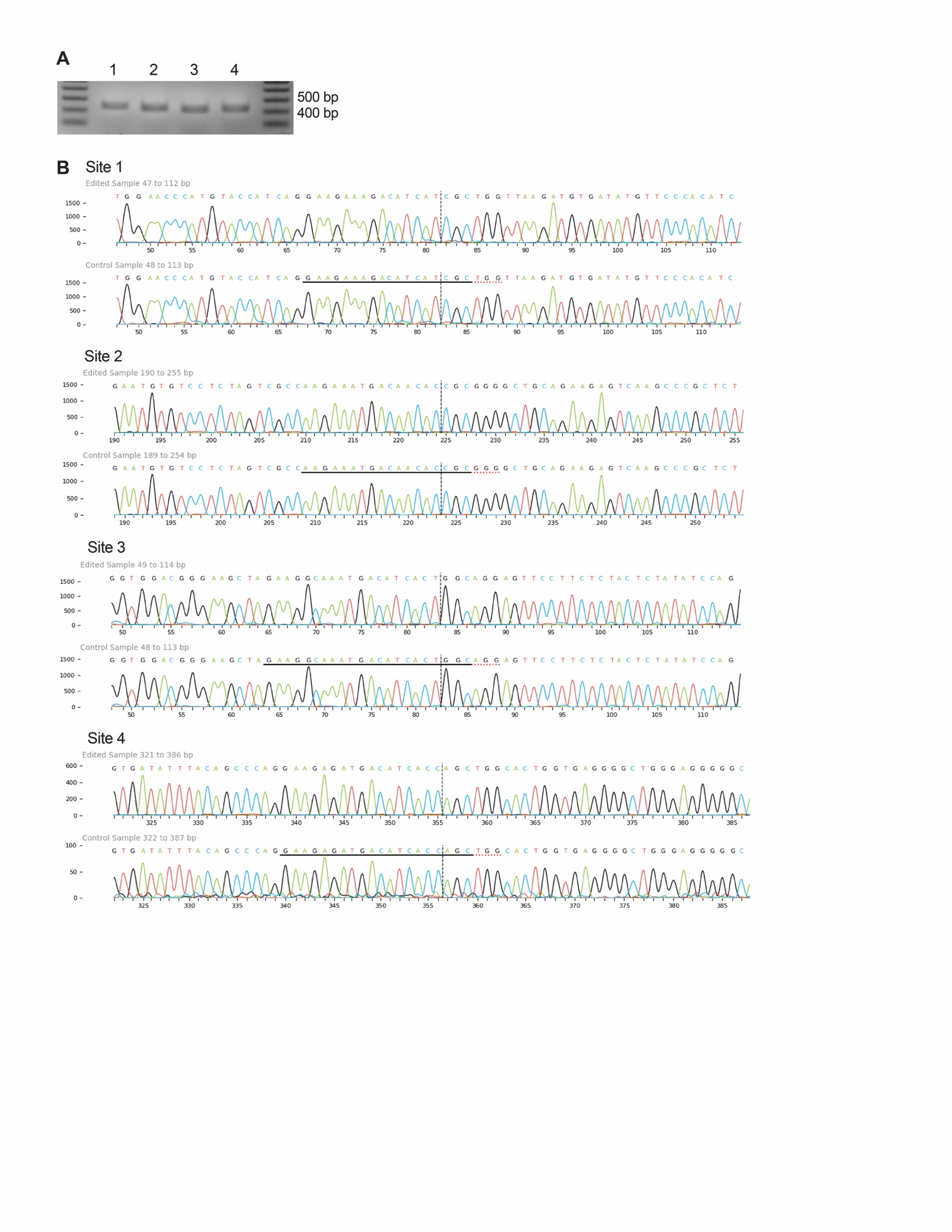

**Supplementary Figure 3. Off-Target analysis of the *CFTR* locus. (A)** Amplicons of the four predicted off-target sites. **(B)** Traces from ICE (Inference of CRISPR Edits) analyses at each site. *n = 3*.

**Supplementary Table 1. Primer and probe sequences**

| **Experiment** | **Primer (5’-3’)** | **Description** |
| --- | --- | --- |
| T7E1 (*CFTR*  locus) | CTCACCTAAGCCTGAGACTAACAAG | forward |
|  | GTGGGTCTTAGAGTACGATTTTCAG | reverse |
| T7E1 (*GGTA1*  locus) | CAGCATCCCTTCCCTCTTCAACTA | forward |
|  | GTATAAAAATCAGGACAATGGCAAC | reverse |
| Junction PCR  (*CFTR* locus) | GTAGGAAGGTGGTTGTCTCTACTTT | LHA forward |
|  | CATAAAACCGACCAAAGAAACTGAC | LHA reverse |
|  | ATCAAGAGGCCAGGACAAGA | LHA sequencing |
|  | AGATCATAATCAGCCATACCACATT | RHA forward |
|  | ATGCCTAAACTTCAGTATCACTTGG | RHA reverse |
|  | TCCATTCAGTGTCAGAGCACA | RHA sequencing |
| Junction PCR  (*GGTA1* locus) | CAGCCCTGGACCTAAATCTTCCTAA | LHA forward |
|  | CTGTTCCGCTCTCTGGAAAGAAAAC | LHA reverse |
|  | AGGAGGGAATGAACAGTGGA | LHA sequencing |
|  | GCATCGCCTTCTATCGCCTTC | RHA forward |
|  | GCAGAGCAAAATGCAGGTCTTATC | RHA reverse |
|  | AAGGGGACAGGGAGACAAGT | RHA sequencing |
| RT-qPCR (*18S*  internal control) | GTAACCCGTTGAACCCCATT | forward |
|  | CCATCCAATCGGTAGTAGCG | reverse |
| RT-qPCR (*K18-CFTR*) | CCTGAGTCCTGTCCTTTCTC | forward |
|  | CGCTGTCTGTATCCTTTCCTC | reverse |
| NHEJ ddPCR (*CFTR* locus) | AGCGAAAAGAAAGGGGAGGT | forward |
|  | CAGCTAGACACCCTCTCACCTG | reverse |
|  | CATCACCGCAGGTCA | VIC probe |
|  | CATCTTCTCCAAACTTT | FAM probe |
| NHEJ ddPCR  (*GGTA1* locus) | ACCATATTCCACTCTGGGTGTATTT | forward |
|  | GGAGACTTTCATCAACATCATTTCA | reverse |
|  | CAGGAGAAAATAATGAATGTC | VIC probe |
|  | CTTGTCTCAACTGTAATGGT | FAM probe |
| ddPCR (*K18-CFTR*) | GATACAGAAGCGTCATCAAAGCA | forward |
|  | GAACTATATTGTCTTTCTCTGCAAACTTG | reverse |
|  | CCAACTAGAAGAGGACATC | probe |
| ddPCR (Ad5) | GCGGTTTTAGGCGGATGTTG | forward |
|  | TCCTCTTATTCAGTTTTCCCGC | reverse |
|  | TAACCGAGTAAGATTTGG | probe |
| Off-target analysis | CTCCCTGTCTGGTTAGCGTC | Site 1, forward |
|  | GATTTCAGTGCCTGCGGAAG | Site 1, reverse |
|  | CAAGGCTGGCATGAAATCTC | Site 2, forward |
|  | CCCCACATGAACCACTGAAA | Site 2, reverse |
|  | ACTCGCCGCTAACCTTTAGT | Site 3, forward |
|  | CATTCTGTTGGCTGTTGCCC | Site 3, reverse |
|  | CAGTCCCCTACTGTCTGGGT | Site 4, forward |
|  | GAGGGACAAGGACAGGTTCTG | Site 5, reverse |
